# Achieving high-quality ddRAD-like reference catalogs for non-model species: the power of overlapping paired-end reads

**DOI:** 10.1101/2020.04.03.024331

**Authors:** Maximilian Driller, Larissa Souza Arantes, Sibelle Torres Vilaça, Tomás Carrasco-Valenzuela, Felix Heeger, Susan Mbedi, Damien Chevallier, Benoit De Thoisy, Camila J Mazzoni

**Affiliations:** Berlin Center for Genomics in Biodiversity Research (BeGenDiv), Berlin, Germany; Evolutionary Genetics Department, Leibniz-Institut für Zoo- und Wildtierforschung (IZW), Berlin, Germany; Universität Potsdam, Potsdam, Brandenburg, Germany; Department Materials and Environment, Federal Institute for Material Research and Testing (BAM), Berlin, Germany; Museum für Naturkunde Berlin (MfN), Berlin, Germany; Université de Strasbourg, CNRS, Strasbourg, France; Institut Pasteur de la Guyane, Cayenne, French Guiana, France; Kwata NGO, Cayenne, French Guiana, France

**Keywords:** ddRAD, single copy orthologs, non-model species, sea turtles, high-throughput sequencing, RAD pipeline

## Abstract

Reduced representation libraries (RRS) allow large scale studies on non-model species to be performed without the need for a reference genome, by building a pseudo-reference locus catalog directly from the data. However, using closely-related high-quality genomes can help maximize nucleotide variation identified from RRS libraries. While chromosome-level genomes remain unavailable for most species, researchers can still invest in building high-quality and project-specific *de novo* locus catalogs. Among methods that use restriction enzymes (RADSeq), those including fragment size selection to help obtain the desired number of loci - such as double-digest RAD (ddRAD) - are highly flexible but can present important technical issues. Inconsistent size selection reproducibility across libraries and variable coverage across fragment lengths can affect genotyping confidence, number of identified single nucleotide polymorphisms (SNPs), and quality and completeness of the *de novo* reference catalog. We have developed a strategy to optimize locus catalog building from ddRAD-like data by sequencing overlapping reads that recreate original fragments and add information about coverage per fragment size. Further *in silico* size selection and digestion steps limit the filtered dataset to well-covered sets of loci and identity thresholds are estimated based on sequence pairwise comparisons. We have developed a full workflow that identifies a set of reduced-representation single-copy orthologs (R2SCOs) for any given species and that includes estimating and evaluating allelic variation in comparison with SNP calling results. We also show how to use our concept in an established RADSeq pipeline - Stacks - and confirm that our approach increases average coverage and number of SNPs called per locus in the final catalog. We have demonstrated our full workflow using newly generated data from five sea turtle species and provided further proof-of-principle using published hybrid sea turtle and primate datasets. Finally, we showed that a project-specific set of R2SCOs perform better than a draft genome as a reference.

## 1. INTRODUCTION

Reduced representation sequencing (RRS) has become very popular in the last several years among scientists studying the genetic variation of non-model organisms (Andrews, Good, Miller, Luikart, & Hohenlohe, 2016; Beichman, Huerta-Sanchez, & Lohmueller, 2018; Narum, Buerkle, Davey, Miller, & Hohenlohe, 2013), with special emphasis on digestion-based techniques, here commonly referred to as RADSeq (restriction-site-associated DNA sequencing). Coupled with high-throughput sequencing technologies, RADSeq libraries produce sequencing data covering thousands to millions of short (<1000 bp) loci throughout the entire genome that can be replicated in large amounts of individuals, within and between related species. The various protocols (reviewed in Andrews et al., 2016) allow for different levels of genome coverage, and are differently suited to studies involving population genetics, phylogenetics, linkage mapping, genomic scans and association mapping. Several of the RADSeq flavours, such as the double-digest RAD (ddRAD, Peterson, Weber, Kay, Fisher, & Hoekstra, 2012) and other similar approaches (e.g. 3RAD, Bayona-Vásquez et al., 2019; and ddGBS Wang et al., 2017), generate fixed-length fragments that must start and end at a recognition site of one of the restriction enzymes utilized. The fixed-length (here generalized as ddRAD-like) protocols offer high flexibility in terms of number of loci targeted and have seen an explosion of publications using the various methods (see Campbell, Brunet, Dupuis, & Sperling, 2018 for an etymological analysis of publications involving reduced-representation methods).

The high flexibility of ddRAD-like protocols relies on the combination of the utilized pair of enzymes and the fragment size selection. Once enzymes have been chosen, researchers can still opt for a specific number of loci, as the range of fragment lengths selected will greatly dictate the extension of genome coverage. However, not only the number of loci will vary, but the complete locus repertoire will change across different ranges of fragment lengths. It has been shown that the set of loci covered can be affected by slight changes in library building protocols (DaCosta & Sorenson, 2014) or by simply including individual libraries in different pools during size selection (Franchini, Monné Parera, Kautt, & Meyer, 2017). Large sensitivity to changes in the protocol affects reproducibility and may not only increase costs but also prevent entire datasets from being compared. Although a few approaches have been suggested in attempts to homogenize the outcome of library building across many specimens (e.g. Franchini et al., 2017), the type of sequencing data frequently generated -such as single-end reads or non-overlapping paired-end reads - and the lack of a high-quality reference genome prevent researchers from understanding what issues may have caused a drop in individual coverage and/or in the set of loci overlapping among samples.

Perhaps the most attractive feature of ddRAD is the fact that no reference genome is needed to perform analyses, as a reference locus catalog can be built *de novo* directly from the data. However, using a reference genome can help improve single nucleotide polymorphism (SNP) calling, depending on the genetic distance to the species analyzed (Paris, Stevens, & Catchen, 2017; Shafer et al., 2017). Even if references can theoretically help SNP calling, draft genomes are prone to incompleteness and high levels of mis-assemblies due to -among other reasons -haplotype divergence in homologous regions (Guan et al., 2020) and extensive repetitive regions (Phillippy, 2017). Moreover, it is still largely unknown how genome assembly quality can affect RADSeq data analysis.

Regardless of whether a *de novo* or reference-based approach is chosen to analyse RADSeq data, the main goal is to ensure that variation among individuals is retrieved from comparisons among orthologous loci (or true homologues), in contrast to paralogous loci. A key step to identify true homologues in *de novo* approaches is the definition of identity thresholds. Alleles with sequencing similarity above an established threshold will be considered as belonging to the same orthologous locus, while more divergent alleles are assumed to belong to different positions in the genome. If thresholds are too high, they will over-split orthologous alleles, whereas too low thresholds will cluster paralogous alleles into the same putative locus. The threshold-definition step is therefore considered to be crucial in *de novo* protocols (Ilut, Nydam, & Hare, 2014) and should be established through an empirically‐ justifiable protocol (McCartney-Melstad, Gidiş, & Shaffer, 2019). If loci are mis-identified, the over- or under-clustering of alleles will affect observed heterozygosity and consequently the results of downstream analyses that depend on genotype frequencies. A common approach to define optimal sets of thresholds (Paris et al., 2017; Rochette & Catchen, 2017) consists of evaluating the changes (i.e. increase/decrease) in numbers of SNPs and variable loci across a set of threshold values. Using a representative and (in principle) well covered part of the sample set, this approach can increase the number of SNPs called. The final choice of parameters, however, depends on the researcher’s interpretation of the output and does not currently include an evaluation of the distribution of pairwise allele identities within or between individuals.

Bioinformatic pipelines commonly used for ddRAD data analysis include Stacks (Catchen, Hohenlohe, Bassham, Amores, & Cresko, 2013), ipyrad (Eaton & Overcast, 2020) and dDocent (Puritz, Hollenbeck, & Gold, 2014), among several others (LaCava et al., 2020). It has been shown that changing parameters and pipelines may strongly affect downstream analyses (Shafer et al., 2017). However, the choice of an optimal combination of pipeline and parameters remains based mostly (or solely) on the highest possible amounts of loci and variation (i.e. SNPs) they output (Díaz-Arce & Rodríguez-Ezpeleta, 2019; Shafer et al., 2017). In contrast, Shafer et al. (2017) hypothesized that the biggest differences among methods are caused by the crucial locus definition step, which is itself highly affected by identity and coverage thresholds. Based on that, a method that specifically focuses on building high-quality reduced-representation locus catalogs can bring a new perspective to the analysis of non-model species, increasing genotyping confidence and consequently the robustness of downstream analyses.

In this study, we present a simple but effective solution to overcome issues in ddRAD-like data that can be applied for any given species or population. By using overlapping paired-end reads, we allow data to be analyzed using haplotypes that represent the original DNA fragments in its entire extension, producing important information on the fragment length and its associated coverage. We have designed a workflow to build and evaluate a reference locus catalog based on a controlled size range and coverage per length. At the end of the workflow, a set of reduced-representation single-copy orthologs (R2SCOs) is identified as the reliable reference to extract variation from across populations or species. The results were compared against locus catalogs built using Stacks and filtered through newly designed scripts that help increase the comparability of the two approaches. Furthermore, we have also developed a new approach to define two identity thresholds for any species based on the generated data. We built R2SCOs independently for each of five different sea turtle species, revealing variable genetic distances and levels of internal heterozygosity. We also used one set of R2SCOs to re-analyze previously published data on hybrid sea turtles in comparison to the available draft genome. Finally, we used published ddRAD datasets including overlapping paired-end reads for three species of new world primates to demonstrate that the approach is easily transferable to other vertebrates.

## 2. METHODS

### Preliminary in silico tests and reference genome

A draft genome of *Chelonia mydas* (CheMyd_1.0, GenBank accession number GCA_000344595.1; (Wang et al., 2013) was used to perform preliminary estimations of fragment yield for different enzyme combinations and size ranges before designing the study, using a dedicated python script (*RAD_digestion*.*py*) that performs *in silico* digestions according to the ddRAD workflow (Peterson et al., 2012). We have chosen the enzyme pair EcoRI (6-bp cutter) and MseI (4-bp cutter) based on the number of expected genome fragments (20,000-30,000 loci) within a size range of 400-500 bp (considered optimal for Illumina sequencing). We have also verified that similar locus counts per fragment size can be found in further related taxa (data not shown). For that, we have digested the genomes of the freshwater turtle *Chrysemys picta* (Schneider, 1783; Shaffer et al., 2013; Chrysemys_picta_bellii-3.0.3, NC_024218.1) and the Chinese softshell turtle *Pelodiscus sinensis* (Wiegmann, 1835; PelSin_1.0, GCA_000230535.1; Wang et al., 2013).

For analyses performed after the sequencing data were obtained, we utilized the version of CheMyd_1.0 genome scaffolded by the DNAzoo project (Dudchenko et al., 2017), identified here as CheMyd_1.0_DNAzoo.

### Library preparation

DNA extractions were performed using the DNeasy Blood and Tissue kit (Qiagen). The preparation of the ddRAD libraries followed the protocol of Peterson et al. (2012) with modifications of adapter sequences according to Meyer and Kircher (2010). In brief, 1 µg of genomic DNA (20-100 ng/µl) was digested with EcoRI and MseI at 37 °C for at least two hours. The ligation of adapters was performed immediately after the digestion. The P5 adapter had inline barcodes of varying sizes (5 to 9 bp) at the restriction site positions and the P7 adapter was designed using one of the strands lacking the complementary region of the indexing primer. Ligated samples were cleaned with 1.8X CleanPCR magnetic beads from CleanNA. A ten-cycle indexing PCR was performed independently for each individual using one of the 50 indexes described by Meyer and Kircher (2010) at the P7 adapter. The indexing PCR was cleaned with 0.8X CleanPCR magnetic beads, and the concentration was measured with Qubit 2.0 using the dsDNA HS assay (Life Technologies) and checked using the Bioanalyzer (Agilent). The indexed libraries were equimolarly pooled before the size selection step.

### Size selection and sequencing

The size selection step was performed using the BluePippin with a 1.5% agarose cassette (250-1,500 bp) and R2 marker (Sage Science). The initial sequencing runs showed a shift in the size selection among different pools and runs (Fig. S1), and therefore the size range selection was extended for the two last runs to ensure overlapping. Libraries were included in four different MiSeq runs and performed with size selections of 495-605 (Run1), 490-610 (Run2) and 450-650 (Run3 and Run4), in order to ensure substantial overlapping across runs. Between 129 bp and 134 bp represented adapter sequences, which means that the longest internal fragments should reach ∼520 bp. Pool amplification was avoided after size selection, and only performed in one run (Run2) with fewer than 10 cycles and in five independent reactions per library, which were pooled and cleaned. The final libraries were characterized with a qPCR using the KAPA SYBR® FAST (Kapa Biosystems) kit and also checked with the Bioanalyzer. The libraries were run on the in-house Illumina MiSeq using the 600-cycle v3 kit.

### Preliminary contamination and homology analysis

Samples were demultiplexed into pools using the Illumina MiSeq Reporter (v2.6.2.1), based on the P7 index. The pools were then demultiplexed into individual samples using the P5 inline barcodes, with Flexbar v.3.0.3 (Roehr, Dieterich, & Reinert, 2017) allowing no mismatches (parameters: -be LEFT_TAIL -u 3). In order to run a preliminary test for major contaminants, a subsample of 50,000 paired-end reads from each individual library was compared against two turtle genomes. For this purpose, the paired-end reads were first mapped against CheMyd_1.0_DNAzoo using bowtie2 v.2.3.0 (Langmead, Trapnell, Pop, & Salzberg, 2009) with parameters adjusted to reach a minimum of ∼80% identity (parameters: --mp 10 --score-min L,-1,-2.0 --no-unal). Unmapped read pairs were compared against CheMyd_1.0_DNAzoo using blastn as implemented in NCBI’s BLAST+ package (v.2.6.0) with a maximum e-value of 10^−20^ and subsequently against *C. picta* (Shaffer et al., 2013; Chrysemys_picta_bellii-3.0.3, NC_024218.1). Finally, reads that did not produce matches were aligned against the GenBank nucleotide (nt) database using the same parameters. The output from blast was visualized using MEGAN v.6.8.18 (Huson et al., 2018).

### R2SCO pipeline

A general scheme of the R2SCO pipeline is depicted in Figure 1. The first two general steps of the pipeline are described in topics below and consist of *(i)* the reads pre-processing, including paired-end reads merging into single sequences (Fig. S2) and *(ii)* the *in-silico* size selection based on the coverage per fragment size estimated from the merged sequences (Figure 1 steps *i* and *ii*).

**Figure 1:**
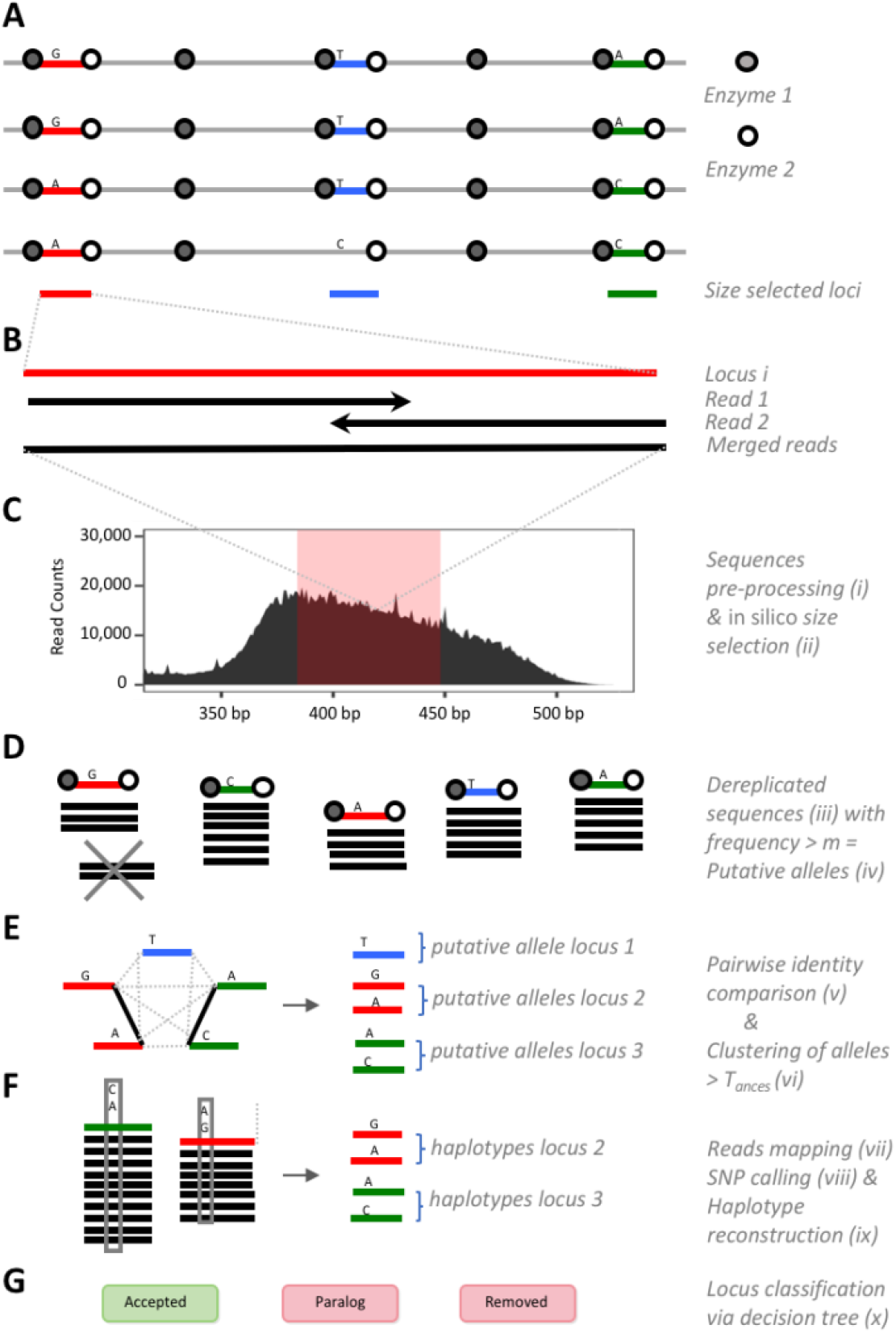
Workflow of the reduced-representation single-copy orthologs (R2SCO) catalog building. The general representation of a ddRAD set of loci sequenced across the genome can be seen in **A**. The reconstruction of an entire locus by merging overlapping reads is shown in **B**. The R2SCO pipeline is illustrated from **C-G** with figures that represent each of the ten steps, from *i* to *x* (identified on the right part of the figure). In **E**, black lines connect putative alleles with identity above *T*_*ances*_ and that will be clustered together, while dotted lines connect alleles presenting pairwise identities below *T*_*ances*_. The decision tree classification workflow is presented in Fig. S3.

Following the steps *i* and *ii*, four other steps represent the definition of loci within and between samples (see Figure 1 and the respective roman numerals): (*iii*) the de-replication (i.e. collapsing of identical sequences) of the preprocessed and size-selected sequences, performed for each individual separately, (*iv*) the definition of putative alleles, also performed by individual, (*v*) the pairwise comparison of putative alleles, performed among all individuals together and finally (*vi*) clustering and definition of putative loci per individual and between conspecific individuals. Steps *iv* and *v* need the thresholds *m* and *T*_*ances*_, respectively, briefly described below:

- *Minimum count per putative allele* (*m*): the minimum coverage per unique sequence (i.e. minimum number of identical sequences after merging reads) within an individual that should represent a putative allele.
- *Ancestral identity threshold* (*T*_*ances*_): the ancestral threshold represents a putative minimum identity above which recent paralogous loci would still align to each other. It can be based on the distance between the most distant species in the analysis or in case of an intra-specific analysis, it can be a conservative value (e.g. 90%).

By using the set of loci generated within each individual, the next four steps of the pipeline add a classification to each locus for every individual independently (see Figure 1 and the respective roman numerals). First, in step *vii* all preprocessed merged sequences are mapped back to the loci (in a two-step-mapping approach, see Supporting Information) of that given individual. For the latter, the merged sequences used must fall within the selected size range with a ±5% extension, to allow the recognition of alleles containing short indels. Subsequently, in step *viii* a SNP calling is performed for each locus individually and in step *ix* the called SNPs are used to “reconstruct” haplotypes.

The last step of the pipeline *(x)* consists of getting all information obtained so far, including putative alleles, mapping results, reconstructed haplotypes and running through a decision tree. Three further thresholds are needed in this step:

- *Minimum locus coverage* (*COV*_*min*_): the minimum coverage per locus within an individual to yield reliable genotypes. It is not used for defining R2SCOs since a locus can be defined by a single allele (i.e. even if the second allele has not been covered, a locus is recognized).
- *Maximum locus coverage* (*COV*_*max*_): the limit between the distribution of sequence coverages of single-copy loci and outlier coverages that mostly represent paralogs.
- *Intra-specific identity threshold* (*T*_*intra*_): the identity threshold within species represents an identity value among sequences that clusters the great majority (at least 95%) of orthologous alleles and excludes the great majority of paralogs. This threshold is obtained by comparing identities among putative alleles within and between individuals from the same species (Supporting Information).

The decision tree (Fig. S3) is run for each individual independently and the output can be roughly subdivided into three types of classifications: *putative paralogs, removed loci* and *accepted loci*. Loci are mostly removed due to incongruencies between the clustered putative alleles and the reconstructed haplotypes from SNP calling. For performing locus classification, the decision tree will utilize: the identity and coverage thresholds defined above (*T*_*intra*_, *T*_*ances*_, *COV*_*min*_, *COV*_*max*_); the evaluation of the remapping of merged reads against each locus; and the comparison between variant positions derived from the putative alleles and from the SNP calling. More details about the decision tree and thresholds can be found in the Supporting Information.

### Sequence preprocessing – R2SCO pipeline step i

The first step performed with the fastq data was the phiX control library cleaning, as it was spiked into every run. Each demultiplexed pool was mapped to an *Enterobacteria phage* phiX174 reference genome (NC_001422.1) using bowtie2 with default parameters. All read pairs that mapped concordantly were removed from the sample. Subsequently, PEAR v.0.9.11 (Zhang, Kobert, Flouri, & Stamatakis, 2014) was used to merge (Figure 1B) the paired-end reads (parameters: -v 30 -n 50). The *RAD_digestion*.*py* script was used to redigest the samples according to ddRAD (options: --dd --rad --q) to account for any undigested or chimeric sequences. To remove non-targeted loci derived from star activity of the enzymes, sequences with incorrect restriction sites at either end of the locus (MseI and EcoRI, respectively) were discarded using the script *checkRestrictionSites*.*py*. Finally, sequences with average Phred quality score below 20 were discarded using Trimmomatic v.0.3.6 (Bolger, Lohse, & Usadel, 2014). The read preprocessing workflow is shown in Fig. S2.

### Chimera estimation

Merged sequences including internal restriction sites could represent either undigested sequences due to low efficiency of restriction enzymes or the ligation of independent digested fragments into chimeric sequences during the adapter ligation step. In order to distinguish both cases, fragments generated from *in silico* digested sequences were analyzed in comparison to each other. The details of the workflow are found in the Supporting Information and Fig. S4.

### In silico size selection – R2SCO pipeline step ii

The range for *in silico* size selection was chosen based on the coverage distribution for each individual (Fig. S5). We have selected a range for which most of the ten individuals showed an estimated mean coverage (i.e. read counts divided by the expected number of fragments obtained from the genome digestion) of at least 20x per locus, while avoiding very high peaks of coverage that could represent highly repetitive paralogous loci (Fig. S5, red arrow). The range and the best run per individual were selected concomitantly. *In-silico* size selection was performed using an awk command (Supporting Information).

For details of steps *iii-x* of the R2SCO pipeline, check the Supporting Information.

### Intra (and inter-)specific set of R2SCOs

Once R2SCOs are defined for each individual, the pairwise results from allele clustering within and -only in case of interspecific analysis -between species are used for defining homology between individuals. Homologous loci across individuals were defined by clustering loci that share at least one putative allele between individuals with an identity above *T*_*ances*_. A new type of paralog could be identified, when two or more loci from within one individual ended up in the same cluster when compared within species or between species. The genotypes in comparisons across individuals from within or between species keep the individual-based classification of the decision tree, but loci will be removed in case they are classified as Paralogs at the level of comparison tested or -in case of interspecific analysis -if they are not present in one or more species of the set. If a locus is present and accepted in one of the two individuals but absent in the other, it is still considered as part of the R2SCO set of that given species.

### Comparisons between individual replicates

Whenever the non-selected run (see above-section *in silico* size selection and Fig. S1) of an individual presented enough coverage (>10x in average) for at least part of the range selected for the R2SCO pipeline, it was used as a technical replicate to evaluate genotyping results from the selected run. Loci from the selected run were only evaluated if they were also covered in the non-selected run. The great majority of the genotypes were identical between selected runs and their replicates (AVG=94.92%, SD=1.35%). In order to evaluate the possible causes for different genotype calling, we have evaluated four different loci categories, where: a) both runs called 1 allele, b) both runs called 2 alleles, c) selected runs called 1 allele and non-selected 2 alleles, and d) selected runs called 2 alleles and non-selected 1 allele.

### Reference genome evaluation

Two types of evaluation were performed using the available draft genome CheMyd_1.0_DNAzoo: 1) A simulation of ddRAD library for a direct comparison with the obtained R2SCO catalog for *C. mydas* and 2) Comparison of references for ddRAD-like data, between the R2SCO catalog produced here for *C. caretta* and the draft genome of *C. mydas* (see the section “Published datasets” below).

To compare the locus catalog and locus classification obtained with the R2SCO pipeline against the CheMyd_1.0_DNAzoo, the genome was digested using the *RAD_digestion*.*py* script to simulate a ddRAD locus catalog. Briefly, the digested genome sequences carrying the overhangs for both enzymes (MseI and EcoRI) were size-selected according to the selected range used in this study (see Results). Digested and size-selected genome sequences were compared pairwise to each other using vsearch (Rognes, Flouri, Nichols, Quince, & Mahé, 2016) v.2.8.6 with option --allpairs_global. The identity threshold was defined by *T*_*ances*_ (here 0.90) and pairwise identity results were used to cluster sequences using the script *clusterFromPairs*.*py*, as described above for the empirical data (Figure 1). As CheMyd_1.0_DNAzoo represents a haploid copy of the genome (i.e. nucleotides represent just one of the chromosomes in the diploid genome), any digested and size-selected sequences forming clusters can be interpreted as ddRAD putative paralogs, as they would represent more than one locus in the genome.

The putative single-copy and paralogous loci were compared with the locus classification obtained through the R2SCO pipeline for the two individuals of *C. mydas* (Cm1 and Cm2). Furthermore, clusters obtained for the genome ddRAD were also used to evaluate the identity among potential paralogs. Note that missing loci in the comparison between R2SCOs and the genome represent mostly mutations in one of the restriction sites (hence changing the size of *in silico* digested sequences) and not the lack of the homologous region in the genome.

### Generating R2SCOs with Stacks

We ran Stacks version 2.53 (Rochette, Rivera‐Colón, & Catchen, 2019) in *de novo* mode using the recommended parameter optimization methods described in Paris et al. (2017) and Rochette and Catchen (2017) for each sea turtle species separately, using all paired-end reads for each selected run with no preliminary filtering. Parameters were tested by fixing *m* to a minimum of three reads, and testing M values between 1 and 8, while fixing n=M. The flag --keep_high_cov was included to remove the stack maximum coverage threshold that is automatically calculated by Stacks based on the entire dataset. A maximum coverage threshold was applied as described below. In order to show the effect of small fragments, we ran Stacks with reads in two sizes: trimmed to 145 bp (simulating 150 bp paired-end reads) and trimmed to 280 bp (original MiSeq 300 bp paired-end reads removing the last base pairs due to a lower quality). Note that read length also determines the minimum locus length that will be analyzed, as shorter reads are discarded.

In order to make the locus catalog produced by Stacks very close to the R2SCO pipeline output, we designed a small workflow (Fig. S6). First, the script *stacksCatalog*.*py* outputs the distribution of mean read coverage and number of loci across the fragment sizes. Subsequently, using the size selection defined by the user, the script *stacks2R2SCOS*.*sh* performs a series of filtering steps directly on the *de novo* locus catalog produced by Stacks for each individual. Filtering consists of the following steps: 1) size selection of loci at a chosen range, 2) removal of loci with internal intact restriction sites, 3) maximum coverage per locus defined as mean + (3*standard deviation) for each fragment size, 4) clustering of loci at 90% and removal of loci with one or more closely-related loci and 5) extraction of singleton loci (clustering with no other locus at 90% identity).

### Statistics tools

Statistical analysis and visualization were performed in Python v.2.7.13 (Rossum & Drake, 1995) using the biopython library v.1.68 (Cock et al., 2009) and in R v.3.6.1 (Team & Others, 2013) using the packages ggplot2 (Wickham, 2016), gplots (Warnes et al., 2015), circlize (Gu, Gu, Eils, Schlesner, & Brors, 2014) and gridExtra (Auguie, Antonov, & Auguie, 2017).

### Sample collection

Tissue samples were obtained for five sea turtle (superfamily Chelonioidea) species: the leatherback turtle *Dermochelys coriacea* (Vandelli, 1761), the green turtle *Chelonia mydas* (Linnaeus, 1758), the olive ridley turtle *Lepidochelys olivacea* (Eschscholtz, 1829), the loggerhead turtle *Caretta caretta* (Linnaeus, 1758) and the hawksbill turtle *Eretmochelys imbricata* (Linnaeus, 1766). Species names are abbreviated as Dc, Cm, Lo, Cc and Ei, respectively. The tribe Carettini is represented by Lo, Cc and Ei, and the family Cheloniidae by all three Carettini species plus Cm. Dc is the only representative of the family Dermochelyidae. For each species, two samples were selected (total n=10) from southwestern Atlantic nesting females -with one exception -from areas that were previously shown to belong to different genetic pools: *D. coriacea* (Martinique and Espírito Santo State in Brazil; Vargas et al., 2019), *C. mydas* (French Guiana and Fernando de Noronha Archipelago in Brazil; Jensen et al., 2019), *E. imbricata* (Rio Grande do Norte and Bahia States from Brazil; Vilaca et al., 2012), *L. olivacea* (French Guiana and Sergipe State in Brazil; Bowen et al., 1997) and *C. caretta* (Bahia State in Brazil and Rio Grande Elevation feeding area; Arantes, Vilaça, Mazzoni, & Santos, 2020a). Nine out of the ten samples were obtained in nesting beaches and should therefore represent the population of origin (Bowen & Karl, 2007). The only sample coming from a feeding area (*C. caretta* from Rio Grande Elevation) presented a mitochondrial haplotype typical from the Indo-Pacific Ocean (Reis et al., 2010). Samples were collected/transported under SISBIO permit 37499-2. Tissue samples were exported from Brazil under CITES permit 14BR015253/DF and imported into Germany under CITES permit E-03346/14. Samples from French Guiana were transported into Germany under institutional CITES between the Leibniz Institute for Zoo and the Wildlife Research and Institut Pasteur in French Guiana.

### Published datasets

Two types of data were utilized for further tests using published and publicly available datasets: 1) Sea turtle backcrossed hybrids 3RAD dataset (Arantes et al., 2020b), including fourteen hatchlings from (*C. caretta* X *E. imbricata*) hybrid X *C. caretta* crossings (n=5) and pure *C. caretta* crossings (n=9) apart from ten species-representative individuals (five from *C. caretta* and five from *E. imbricata*). The data was generated with the 3RAD protocol using the same two main enzymes as in the R2SCOs (i.e. MseI x EcoRI) and a nested size selection range (390bp-410bp). The 3RAD dataset was originally used to identify the presence and type of hybridization between *C. caretta* and *E. imbricata*. 2) New world primates ddRAD dataset (Valencia, Martins, Ortiz, & Di Fiore, 2018). We used data from three species and two individuals per species (n=6) to construct species-specific sets of R2SCOs, including *Lagothrix lagotricha* (Humboldt, 1812), *Saguinus leucopus* (Günther, 1877) and *Sapajus flavius* (Schreber, 1774).

## 3. RESULTS

### Runs summary and in silico size range

Ten individuals (2 from each of 5 sea turtle species) were sequenced in four different MiSeq runs (Fig. S1). Nine out of the ten individuals were sequenced in replicate, always in different runs. The four runs yielded very different distributions, partially due to the increasing size range selection from subsequent runs (performed to guarantee overlapping across a substantial size range as size selection was hard to reproduce exactly), but also due to strikingly different levels of small fragments largely outside of the selected range for each run (Figure 2 and Fig. S1). The size range chosen for the selected samples was 384-448 bp, avoiding a very high peak observed in all species but *D. coriacea* at sizes 449-450 bp. One individual from *L. olivacea* (Lo2) only reached the desired coverage for part of the selected range (Figure 2).

**Figure 2:**
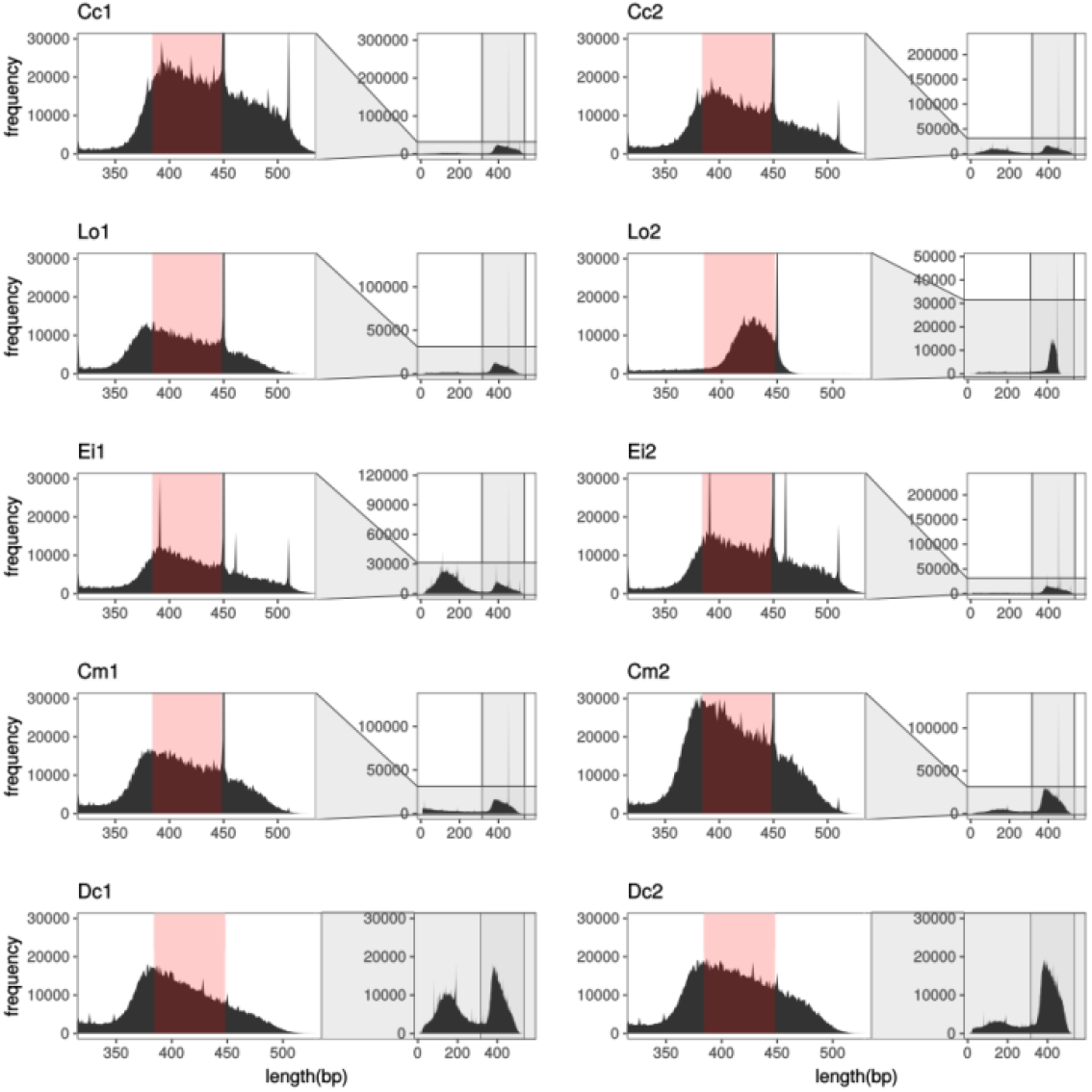
Fragment size distribution of merged and preprocessed reads for 10 sea turtle individuals from 5 species, including only the selected run for each individual. The zoomed-in area on the left side shows the main distribution area for each individual and highlights in light red the *in-silico* selected size range. The plots on the right side for each individual show the entire fragment size distribution, including the very high peaks at size 450 bp.

### Preprocessing

The average number of raw read pairs per individual was 2,968,893 (SD=1,110,900, Table 1). Across all ten selected samples an average of 98.47% (SD=0.17%) of the raw demultiplexed reads remained after all pre-processing steps. For each selected sample, >99% of the reads merged correctly. The *in-silico* digestion showed a frequent occurrence of internal intact restriction sites in the sequences (see *Chimera detection* section below), representing between ∼6% and ∼37% of the merged sequences across individuals (AVG=15.1%, SD=8.7%). All sequences digested *in silico* that presented the correct restriction site combination were kept for further analysis steps, regardless of their new length. Overall, the vast majority of the remaining sequences presented the correct restriction sites on the edges (AVG=99.2%, SD=0.16%) and virtually all the remaining sequences (>99.97%) presented average quality above Phred score 20. Although only a small proportion of reads were filtered out during preprocessing (Table 1), the large amounts of small sequenced fragments – further inflated by *in silico* digested sequences -caused the number of remaining sequences lying within the selected size range (384-448 bp) to drop substantially across individuals. Final preprocessed and size-selected sequences ranged from only ∼13% of the initially demultiplexed reads in Ei1 to ∼57% in Lo2 (AVG=34.03%, SD=11.69%). The statistics for each step can be seen in Table 1 for all sizes and limited to the size range selected.

**Table 1:**
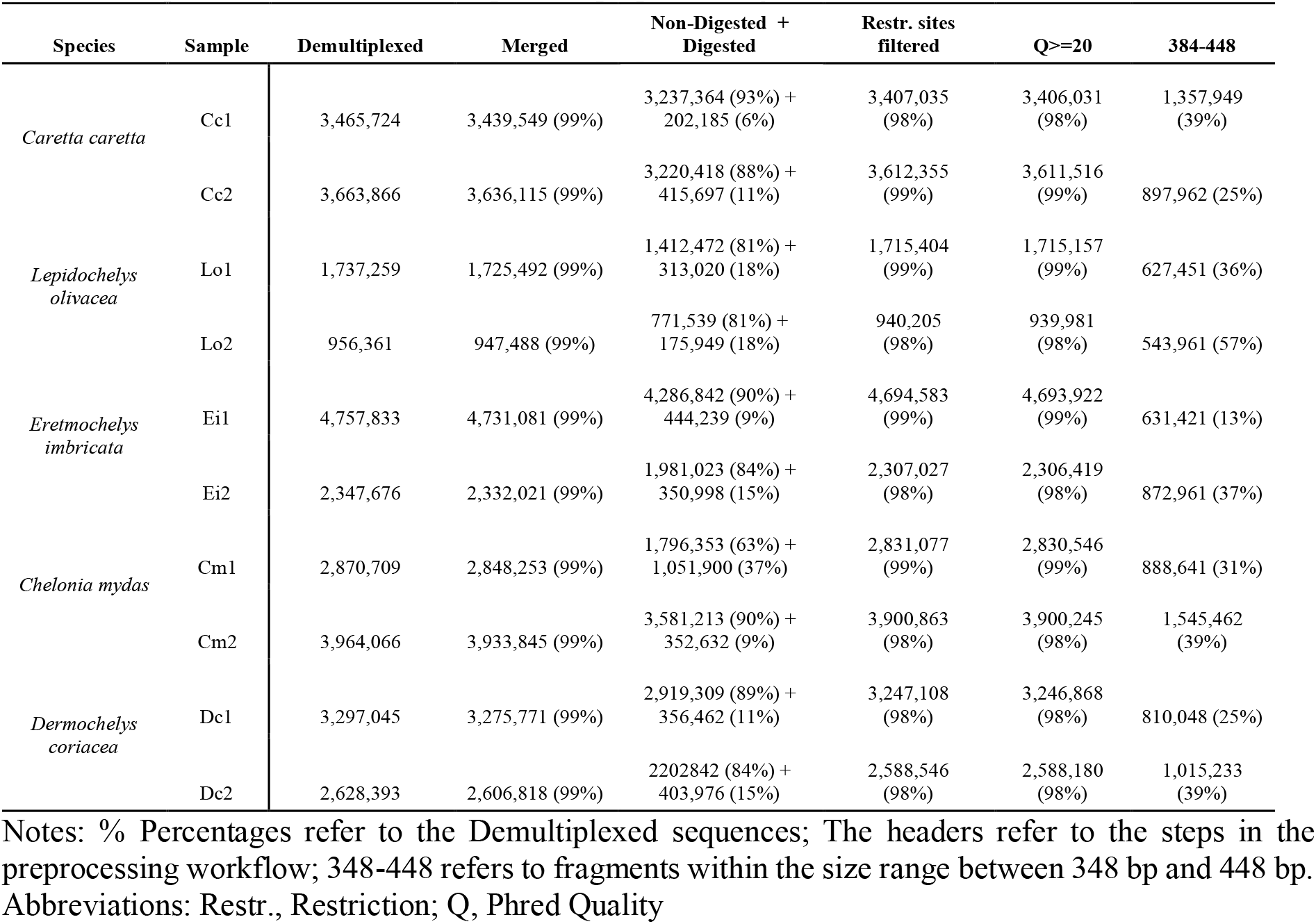
Sequence counts for each step of the preprocessing workflow and after size selection

### Chimera detection

The only significant drops in sequence numbers within the *in silico* selected size range (384-448 bp) during the preprocessing step happened after *in silico* digestion (Fig. S7A). Using our newly developed chimera detection pipeline (Supporting Information and Fig. S4), we were able to differentiate between chimeric sequences and undigested fragments (Table S1). The individual Cm1 presented the highest frequency of sequences with internal restriction sites (∼37%, Fig. S7B). Of the sequences with at least one internal restriction site on average 53% (SD=24.4%) could be attributed to incomplete digestion, while an average of ∼34.6% of the sequences represented a chimeric construction.

### Contamination and turtle homology

The preliminary (optional) contamination analysis performed with a subsample of non-filtered 50,000 paired-end reads per individual revealed very little contamination for each individual (Fig. S8). In contrast, the analysis indicated very high levels of homology among sea turtles. All individuals from the five species presented very high proportions of successful read mapping to the *C. mydas* genome using bowtie2 (between ∼94% and ∼98% of the subsampled reads, Table S2). By adding the two blast comparisons of non-mapped reads against the genomes of *C. mydas* and the western painted turtle *C. picta*, all ten individuals reached homology levels against available turtle genomes above 97%.

### R2SCO pipeline

Putative alleles were defined as sequences within an individual with a minimum coverage of three (m≥3) and sizes between 384 and 448 bp, after the *in-silico* size selection. Putative alleles were compared pairwise within and between all 10 individuals in order to define the two identity thresholds, *T*_*ances*_ and *T*_*intra*_. The distribution of allele identities between *D. coriacea* and each of the other species was used to define a conservative ancestral threshold (*T*_*ances*_=0.90, Figure 3A and 3C). For each species, the pairwise comparison between putative alleles within individuals and between individuals helped establish the intra-specific threshold (T_*intra*_, Figure 3A-B). The *T*_*intra*_ values for Cc, Lo, Ei, Cm and Dc were respectively set to 0.989, 0.991, 0.987, 0.98 and 0.991. *COV*_*max*_ ranged from 75 to 131 mapped reads depending on the individual (Fig. S9).

**Figure 3:**
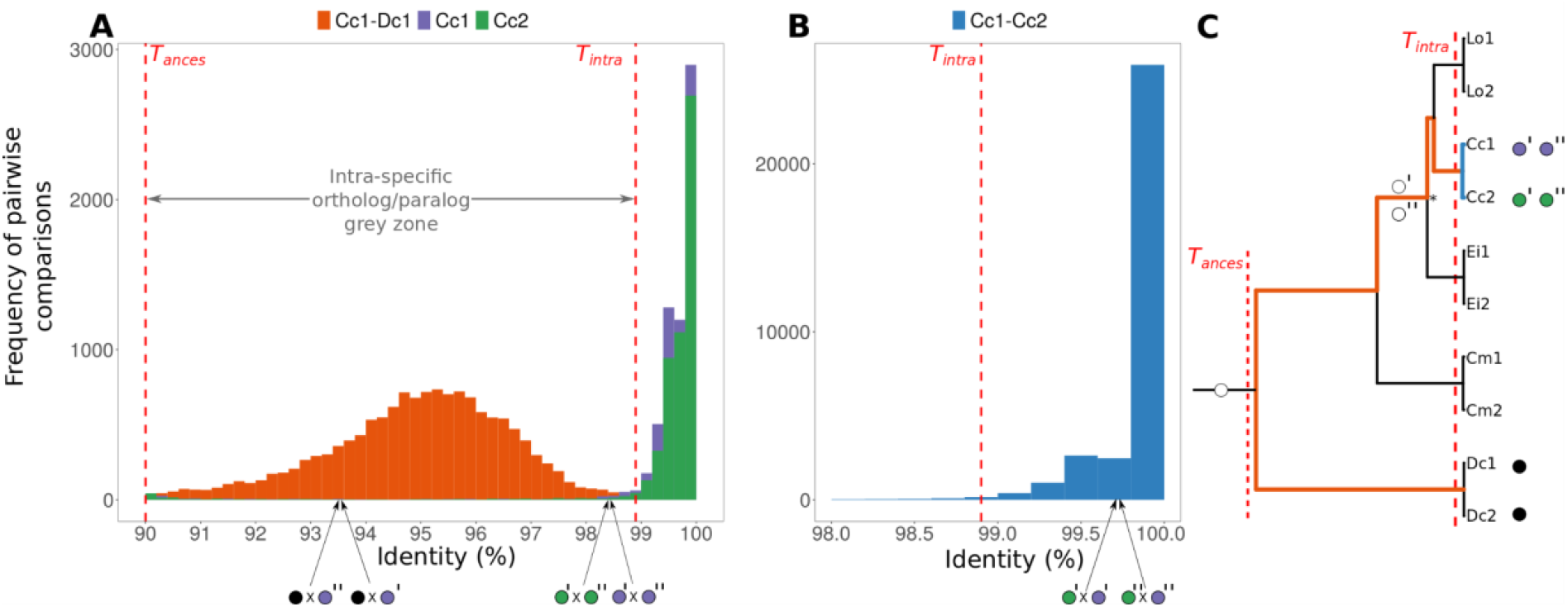
Definition of identity thresholds (*T*_*ances*_ and *T*_*intra*_) based on putative alleles pairwise identity. The figure shows the example of *C. caretta* individuals Cc1 and Cc2 and the comparison with the *D. coriacea* individual Dc1. *T*_*ances*_ and *T*_*intra*_ are shown as dashed red lines. Circles represent alleles from a locus that suffered a duplication where purple, green and black represent Cc1, Cc2 and Dc1-2, respectively. Single and double apostrophes refer to paralogous loci, where orthologs have the same number of apostrophes. In **A**, the whole grey zone between thresholds is supposed to majoritality include identities between putative alleles of paralogous loci. The comparison between individuals in **B** shows the high amounts of identical alleles shared between them. The tree in **C** represents the sea turtle phylogeny and shows a hypothetical duplication of a locus before the tribe Carettini split (node marked with *). The identity relation between alleles from this locus is shown in all 3 figures, exemplifying where identity between paralogs are supposed to fall. Circles indicate an example of a homologous locus for all sea turtles, but with a duplication event within Carettini (indicated in **C**).

The number of accepted R2SCOs within each species was very homogeneous, ranging between 22,919 and 24,227 loci (Figure 4). Rejected loci were more numerous in *C. mydas* (n=5,088) followed by *E. imbricata, C. caretta, L. olivacea* and *D. coriacea*, ranging from 5.32% to 18.17% of the total number of initial loci. Comparing species pairwise within the three main phylogenetic clades (Carettini, Cheloniidae and Chelonioidea) also revealed homogeneous numbers of loci in common, while the total number of common loci for all species from each clade decreased as the last common ancestor became more distant (Figure 4, bars with asterisks). The number of accepted R2SCOs across all species within Carettini, Cheloniidae and Chelonioidea were respectively 14,906, 10,203 and 5,526 (Figure 4).

**Figure 4:**
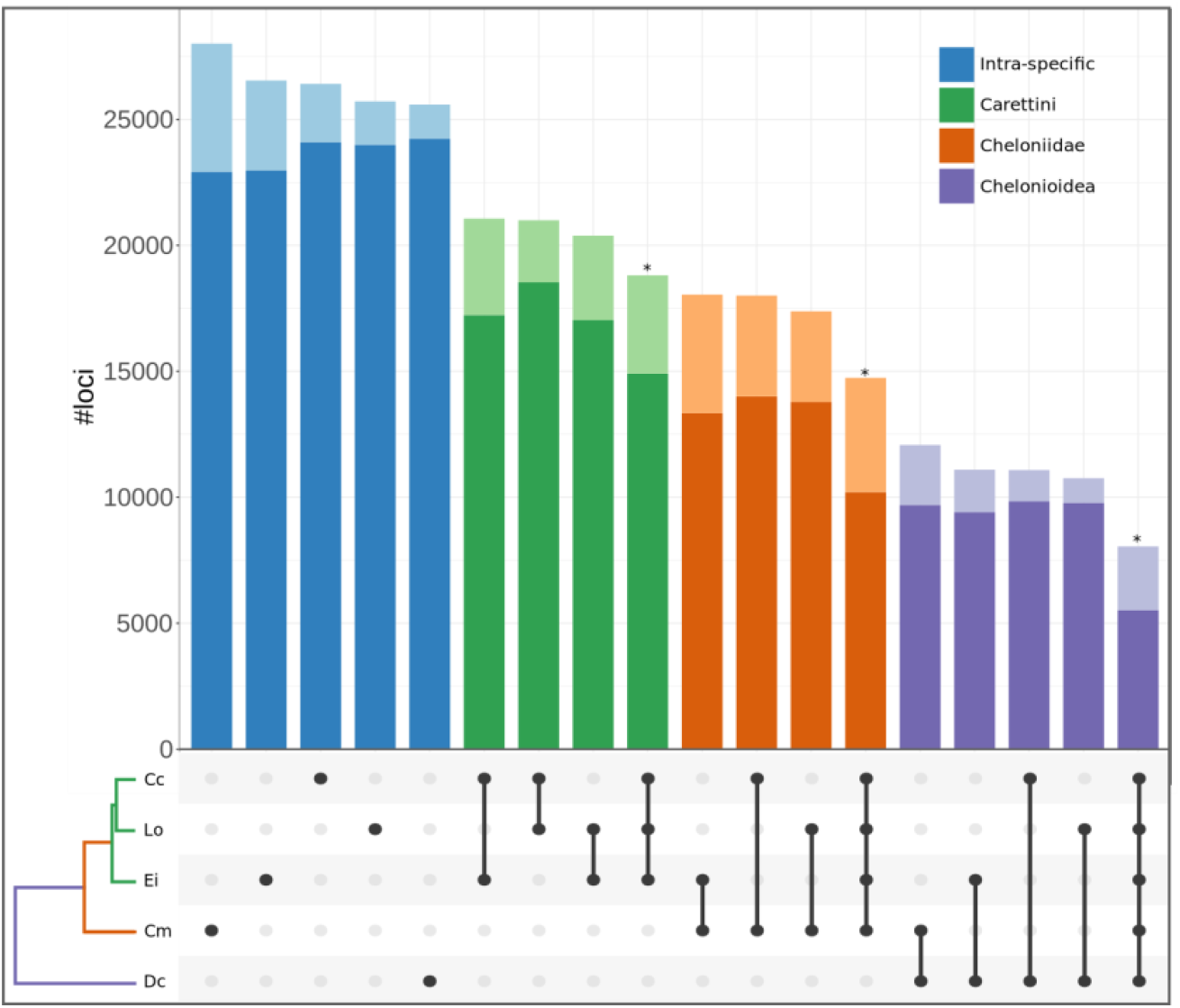
Number of putative loci within and shared between species. Darker shades represent the accepted loci and lighter shades the loci removed by the decision tree. The * denotes comparisons between all species within the corresponding tribe/family/super-family. The lower part of the figure shows which species were tested together, matching the corresponding bar above.

Locus counts across fragment sizes were very similar among all five species as well as the CheMyd_1.0_DNAzoo genome digested *in-silico* (Fig. S10A), including the same trend of decreasing locus numbers at increasing fragment sizes. The latter is also accompanied by a decrease in the number of reads across sizes, generating a slightly decreasing average read count per locus for some of the individuals (Fig. S10B).

### Genotypes and Decision tree evaluation

Six out of the nine non-selected runs were used as replicates in a genotype accuracy analysis. The comparisons were plotted for three individuals (Cc1, Cm2, Ei2) in Figure 5, and include information about allele coverage for both selected and non-selected runs, as well as decision tree classifications for the selected runs (Table S3). For comparing both replicate runs, we have defined putative ‘true genotypes’ based on the following criteria: 1) Selected runs genotypes were only considered true if they got a “Accepted Allele” classification; 2) Homozygous loci were considered true if coverage was higher than 15 reads; 3) Heterozygous loci were considered true if locus coverage was higher than 15 and alleles presented a balanced coverage lying close to the diagonal as seen in Figure 5D, which presents only those loci identified as heterozygotes in both replicates.

**Figure 5:**
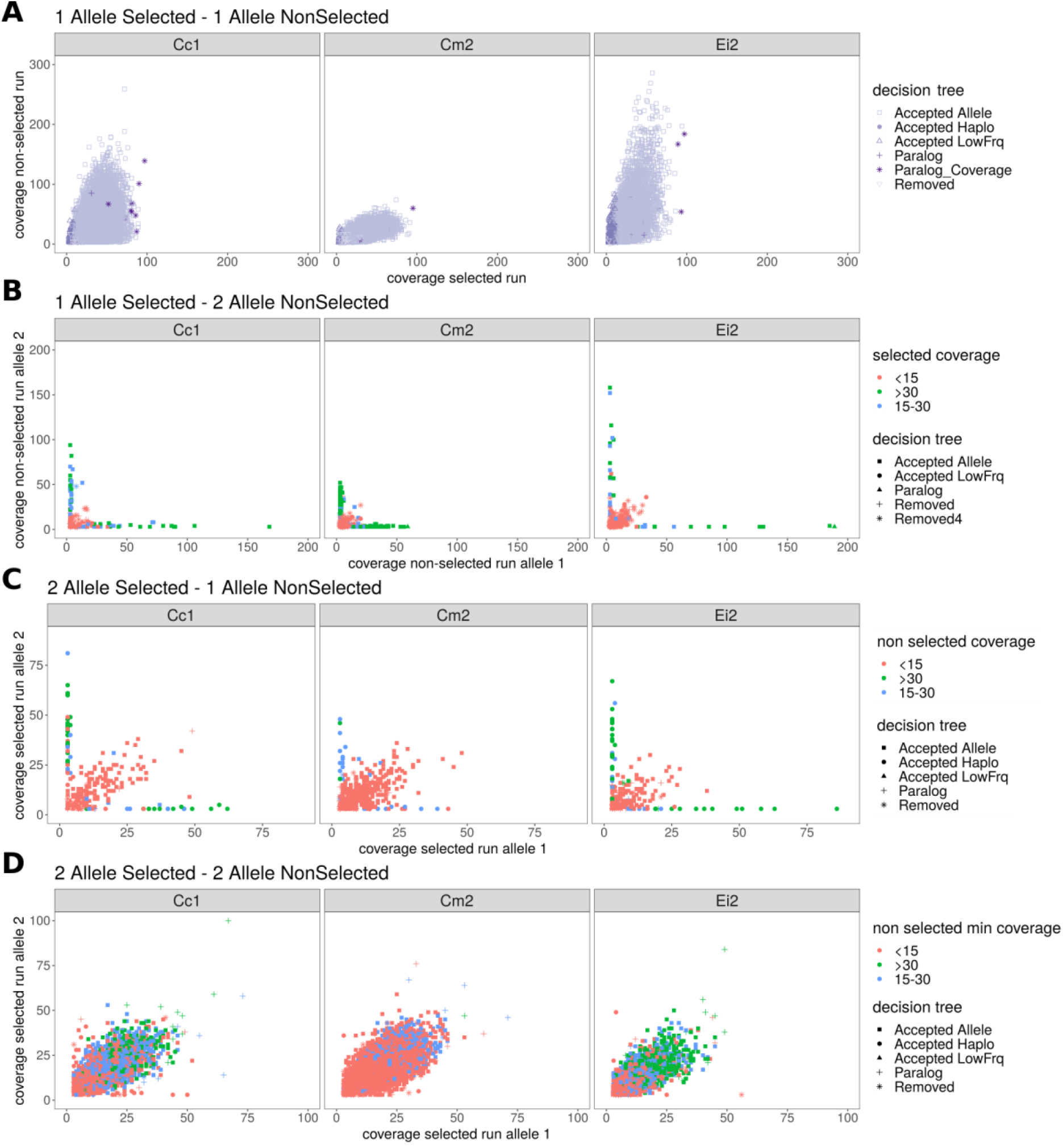
Analysis of replicate runs (selected and non-selected) for three individuals. Comparisons are shown for **A**) loci where both runs agree on one allele, **B**) cases where the selected run has one allele and the non-selected run has two, **C**) the selected run has two alleles and the non-selected one and **D**) where the selected and non-selected run agree on two alleles. Colours on the bottom figure represent the minimum coverage between the two alleles from the non-selected run. All classifications coming from the decision tree represent only selected runs.

Comparisons between replicates show that low coverage really seems to affect the genotyping confidence, while high coverage can bring artifacts to an allele-level coverage. For example, true heterozygotes in the non-selected run that presented a single allele in the selected run (Figure 5B, diagonal) were mostly low coverage (i.e. <15 reads) in the latter and got a “Removed” classification by the decision tree. Similarly, true heterozygotes in the selected run that presented a single allele in the non-selected run (Figure 5C, diagonal) were mostly low coverage (i.e. <15 reads) in the latter. In contrast, false heterozygotes often presented high coverage but a clear unbalance between coverage of both alleles. For instance, we observed that true homozygotes in the non-selected run that presented two putative alleles in the selected run (Figure 5C, two perpendicular lines at low values for one axis) were classified based on SNP calling as homozygous by the decision tree and had very low coverages for the minor putative allele. The false heterozygous presented mostly high coverage (>15) in the selected runs. Likewise, true homozygotes in the selected run that presented two putative alleles in the non-selected run (Figure 5B, two perpendicular lines at low values for one axis), presented very low coverages for the minor allele in the latter.

Finally, one further observation from comparing the replicate runs per individual include the fact that only a few of the loci classified as paralogs due to high coverage in the selected run were also highly covered in the replicate run (Figure 5A).

### Stacks locus catalogs x R2SCOs

We ran Stacks with the default *de novo* mode using thresholds optimized for each individual, and subsequently applied several filters to approach the Stacks catalog to the R2SCO sets (Fig. S6). We performed two Stacks runs keeping all loci generated in the catalog: one with paired-end reads trimmed to 145 bp and one with paired-end reads trimmed to 280bp. The first length included shorter loci (starting in 145 bp) and have generated a much higher amount of loci in comparison to 280 bp (Figure 6), with striking differences occurring in the three individuals -Dc1, Cc2 and Ei1 -with the highest proportion of small fragments (as seen in Figure 2). Differences between the two read lengths were however much lower in numbers of polymorphic loci, partially due to the fact that at 2×280 bp loci are covered in their entire extension (i.e. reads are overlapping throughout the length distribution) and -in contrast to 2×145 bp reads -any SNP present in a locus is detected. Moreover, there was a substantial increase in locus coverage and heterozygosity per bp when excluding small fragments, suggesting that the level of variability detected is highly affected by locus average coverage.

**Figure 6.**
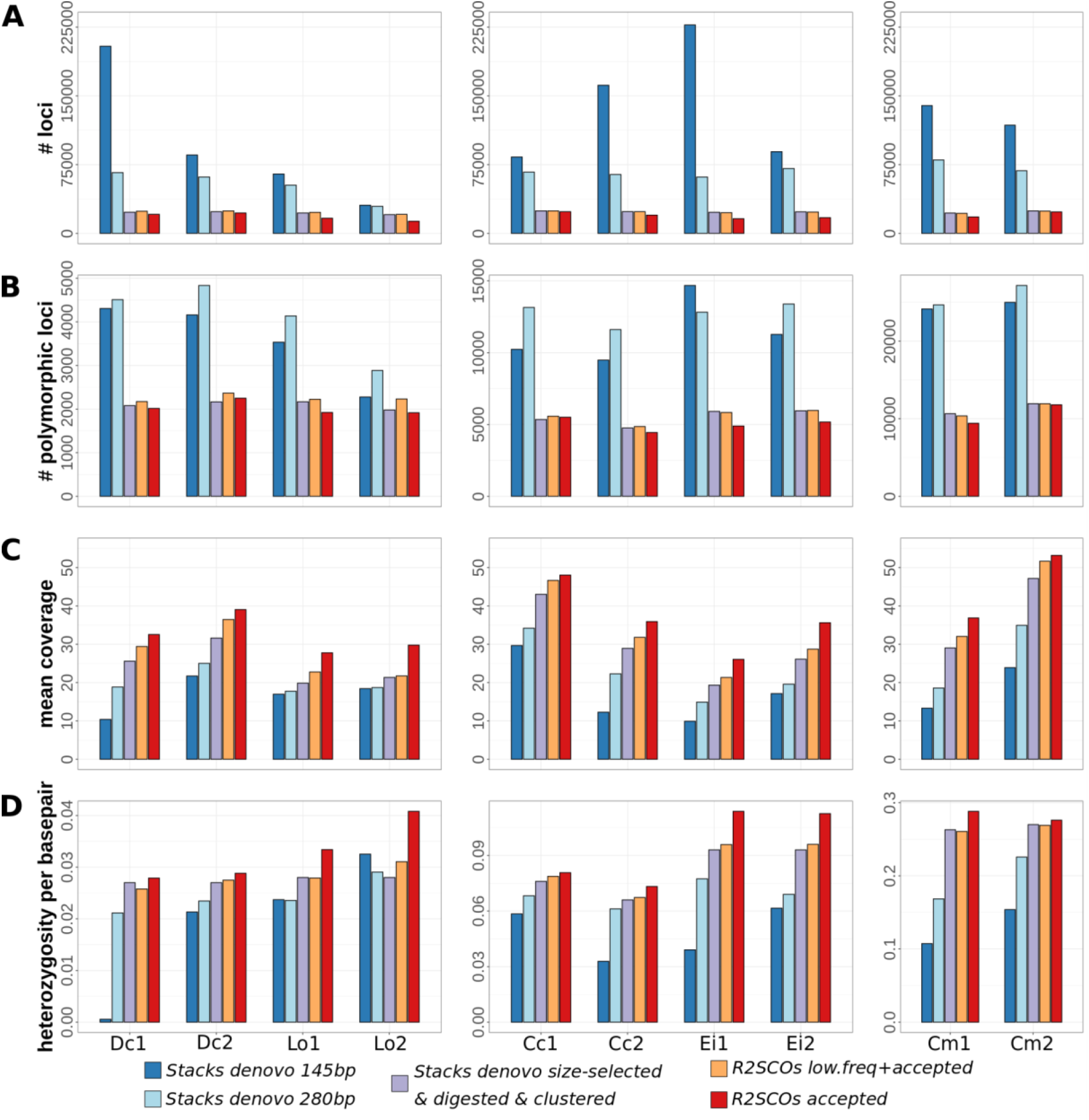
Comparison between R2SCOs and the Stacks *de novo* pipeline using different settings: 1) Stacks *de novo* using reads trimmed to 145 bp, 2) Stacks *de novo* using reads trimmed to 280 bp 3) Stacks *de novo* using reads trimmed to 280 bp filtered to approach R2SCOS: size selected for loci 384-448bp, digested *in silico* and clustered at 90% identity to remove paralogs, 4) R2SCOs including low frequency (< 15 reads) loci and 5) R2SCOs including only loci with coverage >=15. The comparison showcases the A) number of loci obtained in the catalog, B) number of polymorphic loci, C) mean coverage per locus D) heterozygosity per base pair.

When applying R2SCO filters -i.e. restricting the analysis to the selected size range (384-448 bp), removing loci containing internal restriction site and removing paralogs with identity higher than 90% - to the Stacks *de novo* catalog, there is a considerable drop in locus count as well as in the number of polymorphic loci in comparison to the entire dataset using the same 2×280 bp reads (Figure 6A and 6B). However, Stacks locus counts are extremely variable across individuals (between 29,502 and 80,107) in the latter, while they remain stable when the same size range is considered (between 20,410 and 24,750). Furthermore, when applying R2SCO filters there was a substantial increase in the mean coverage per locus (Figure 6C, +6.8 on average, SD: 3.3) and heterozygosity per base pair (Figure 6D, +0.02 on average, SD: 0.03) for most individuals.

When comparing Stacks results using R2SCO filters and the actual R2SCOs, values became very similar across individuals for locus counts, number of polymorphic loci and polymorphism level per base pair (Figure 6A-B,D). Interestingly, the average coverage for Stacks-with-R2SCO-filters loci is consistently lower in comparison to R2SCOs (Figure 6C). Finally, when only R2SCOs with coverage above 15 were considered, both the levels of heterozygosity and coverage per locus increased significantly (+0.01 and +7.53 averaged across all species), indicating coverages below 15 cause an important inflation of homozygosity. Significant changes in coverage and heterozygosity per locus between the different sets of loci were tested for using an one sided Wilcoxon test in R (*α* = 0.01).

Subsequently, we compared the overlapping between R2SCOs and Stacks-with-R2SCO-filters locus catalogs for each species (Figure 7A for Cm and Table S4 for the other four species). Between 89.75% and 94.32% of the combined number of loci were identified by both pipelines. Loci accepted by both pipelines ranged from 77.84% to 90.74% of the total loci. Note that loci rejected by Stacks during the first filtering steps are not present in the output and can therefore not be analyzed. Stacks missed the highest levels of loci (2420 to 3221 loci with any R2SCO classification and between 1417 to 1949 of accepted R2SCOs, depending on the individual). In contrast, very few loci (between 299 and 382, per individual) accepted by Stacks were missed by the R2SCO pipeline.

**Figure 7.**
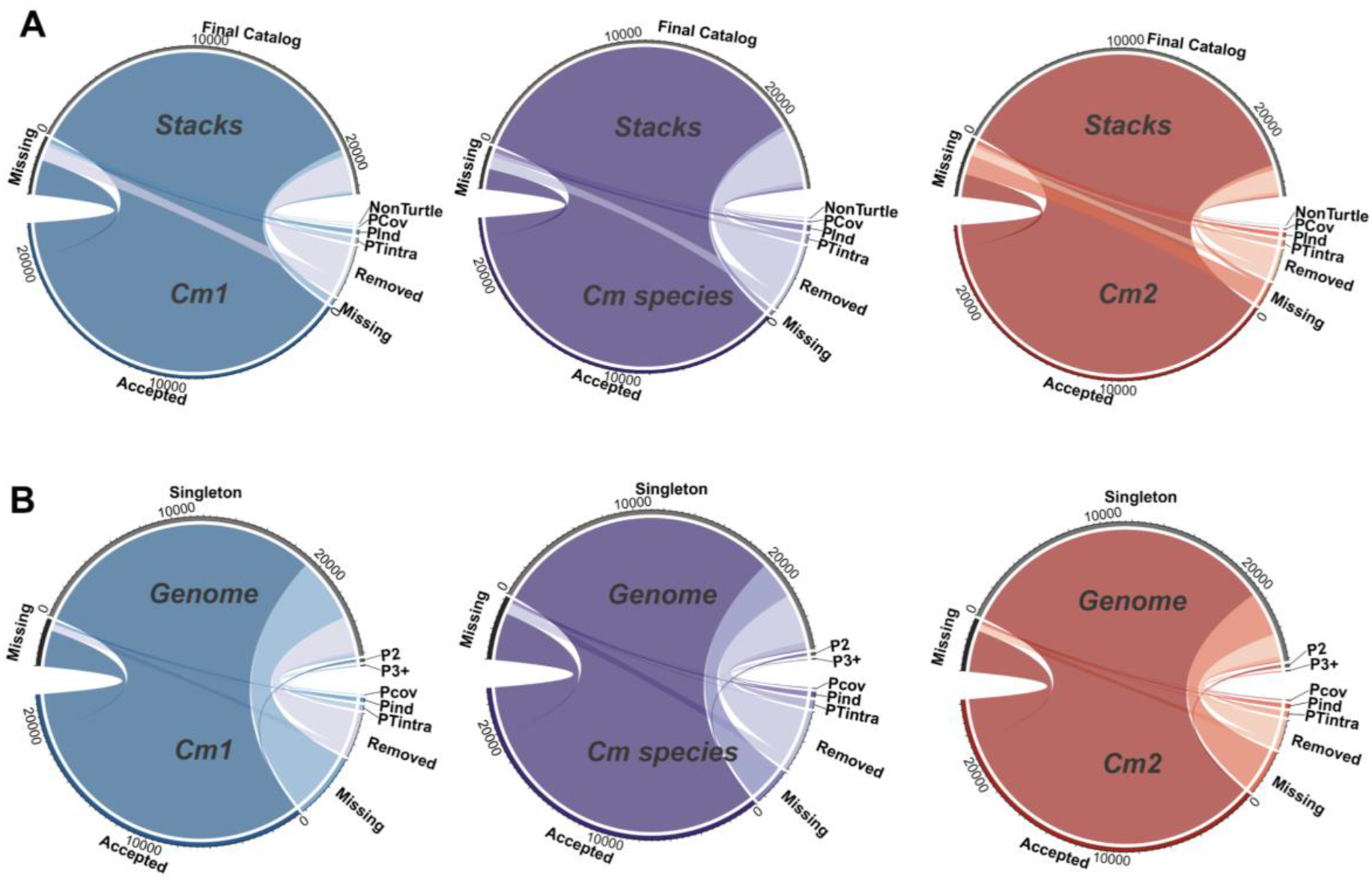
Circos plots showing comparisons of two *C. mydas* individuals (Cm1 and Cm2) and their combined loci (Cm species) against the Stacks final catalog (**A**) and genome clusters of digested sequences (**B**). In (**A)**, the Final Catalog refers to Stacks locus catalog filtered with the Stacks2RSCOs workflow. For the genome (**B**), loci are either missing (present in the individuals but not in the genome), singleton (cluster of one locus), P2 (clusters with 2 loci) or P3+ (clusters with three or more loci). For the Cm individuals, loci can be accepted (as single copies), missing (if present only in the genome digestion), removed by the decision tree or classified as paralogs based on coverage (Pcov), *T*_*intra*_ (PTintra) or the presence of more than 2 alleles (Pind).

### Reference Genome x R2SCOs

We have compared the results from the two *C. mydas* individuals independently (Cm1 and Cm2) and in combination (Cm species) against the CheMyd_1.0_DNAzoo genome after *in silico* digestion with and without a clustering step (Figure 7B). The great majority of accepted loci in the two individuals were found as singletons on the genome (88.47% for Cm1, 87.99% for Cm2 and 85.43% of the combined Cm species). The analysis shows that paralogy is rare within the size range we selected *in silico* (Figure 7B), reaching only 1.04% of the genome digested fragments. In contrast, missing loci between the digested genome and the two individuals reach up to ∼16% of the genome loci, ranging from 1,212 to 4,229 loci, depending on the comparison (Table S5). Similarly, there is a high proportion of removed loci for Cm1 and Cm2 (11.28% and 7.23%, respectively), most of which were found as singletons in the genome. Removed loci within individuals are clusters that -although not identified as paralogous within the first steps of the decision tree -were removed due to at least one type of genotyping issue identified further down in the decision tree. A more detailed analysis of the comparison between Cm individuals and the genome can be found in the Supporting Information.

### Sea turtle genetic variability

In order to compare the genetic variability across sea turtle species, we used the set of 5,526 R2SCOs for the whole superfamily Chelonioidea (Figure 4), excluding the accepted genotype low frequency (i.e. with <15 mapped reads) category for each individual. The numbers of loci used in this analysis for each individual can be seen in Table S6.

The levels of heterozygosity found for conspecific individuals were very similar, but diverged greatly among species (Figure 8A). *Chelonia mydas* was the most variable species, almost three-fold higher than the second most heterozygous one, *E. imbricata. Caretta caretta* individuals were the third most variable and presented very similar heterozygosity values despite deriving from very distant locations (Atlantic vs. Indo-Pacific). *Lepidochelys olivacea* and *D. coriacea* both presented extremely low levels of heterozygosity (<0.005%), roughly 10-fold smaller than *C. mydas*. Among the five sea turtle species evaluated in this study, *D. coriacea* seems to present the lowest genetic variability.

**Figure 8.**
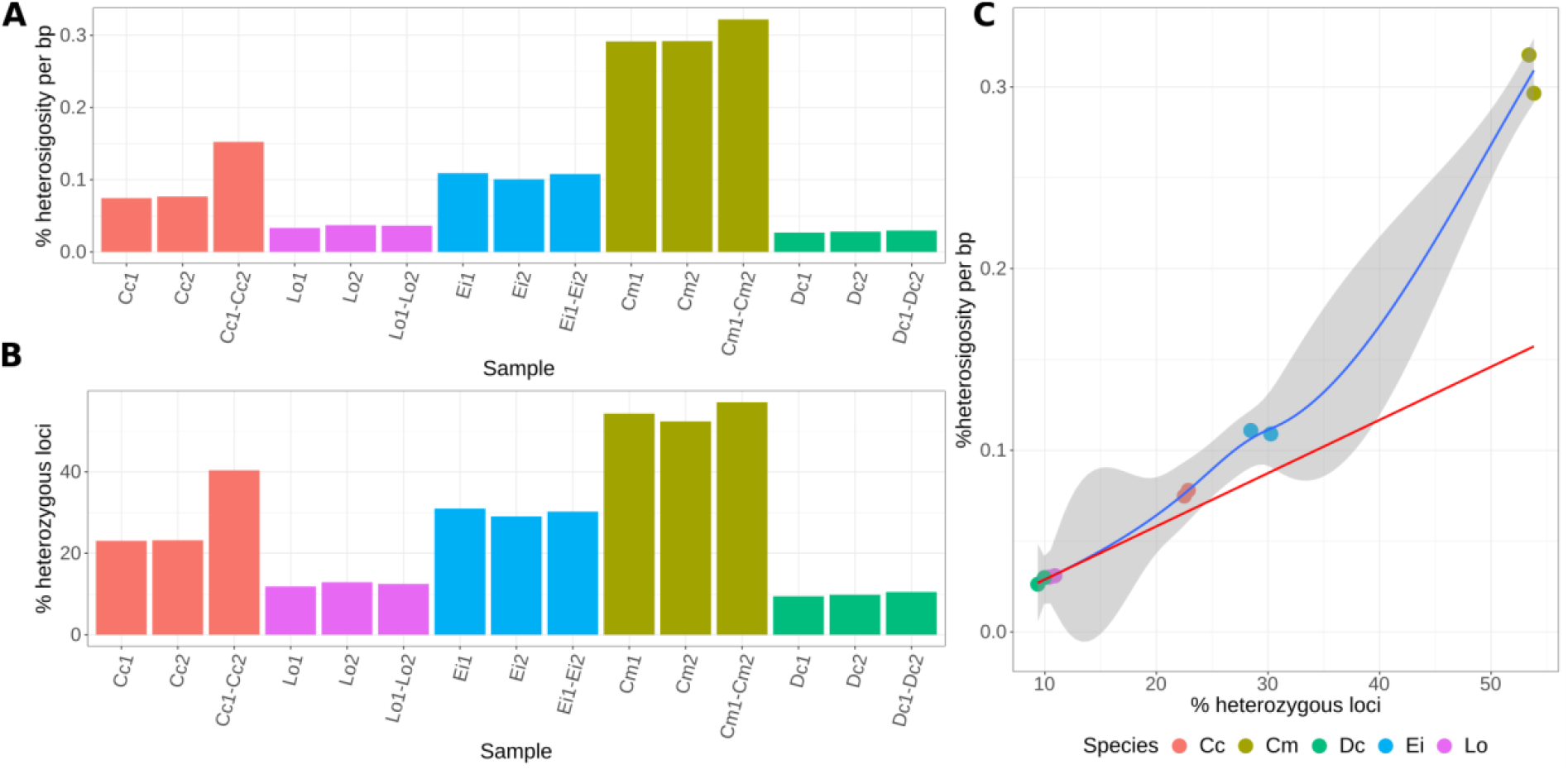
Heterozygosity levels at Chelonioidea R2SCOs. In **A** and **B**, the third bar for each species corresponds to the analysis of conspecific individuals. In **A**, the heterozygosity was estimated for each individual and genetic distances between individuals. **B** shows the proportion of variable loci for each individual and the combining results from two individuals. In **C**, the relation between heterozygosity per base pair vs. the proportion of heterozygous loci is shown for each individual. The blue line is a loess regression based on all values and the red line is a linear regression based on values for Dc and Lo. The grey area around the blue line highlights the 95% confidence interval of the loess regression based model.

The third histogram bar for each species in Figure 8A shows a comparison of pairwise genetic distances between individuals. For all four species for which both individuals belong to southwestern Atlantic populations, there was only a slight increase in the level of variability when comparing both individuals in relation to intra-individual analysis, indicating that both populations share a major part of their genetic variability. This is however not true for *C. caretta*, for which individuals originated from different ocean basins and seem to have quite divergent alleles, reaching twice as much the distance between alleles found within individuals (Figure 8A, Cc1-Cc2 bar).

The amounts of variable loci per individual follow the trend of species heterozygosity levels (Figure 8B), with *C. mydas* individuals presenting the highest proportion of heterozygous loci (AVG=53.61%), while *L. olivacea* and *D. coriacea* present roughly only 10% of variable loci within individuals. However, there is a clear tendency towards saturation as the average number of SNPs per locus increases (see the theoretical red line in Figure 8C based on *L. olivacea* and *D. coriacea* values). Finally, in order to check if the heterozygosity levels calculated based on the R2SCOs at the Chelonioidea level could be biased due to the conservation of restriction sites across all five species, we estimated heterozygosity at common locus sets for different levels of phylogenetic distance within and among species. Very similar values of heterozygosity were found across the different levels of R2SCOs (Fig. S11).

Using the 5,526 Chelonioidea R2SCOs to estimate pairwise genetic distances among all individuals, we obtained the true phylogenetic relationships among the five sea turtle species, as seen in a heatmap analysis (Fig. S12). *Dermochelys coriacea* showed an average of 4.67% (SD=0.11%) genetic distance against the other four lineages, while showing slightly smaller distances against *C. mydas* (AVG=4.48%). *Chelonia mydas* presented very similar genetic distances (2.32%, 2.39% and 2.35%) against each of the three species from the Carettini tribe. *Lepidochelys olivacea* and *C. caretta* presented the smallest genetic distances (AVG=0.96%), while *C. caretta* had a slightly smaller genetic distance against *E. imbricata* compared to *L. olivacea* (averages 1.08% and 1.15%, respectively). The genetic distance matrix among all individuals from the five species can be found in Table S7.

### Sea turtle hybrids

3RAD data from 14 sea turtle hatchlings sampled from six nests with suspected hybridization (Arantes et al., 2020b) were re-tested. Paired-end non-overlapping reads were mapped against three different references: 1) the CheMyd_1.0_DNAzoo genome without any treatment, 2) CheMyd_1.0_DNAzoo digested and size-selected based on a well-covered size range for the study (390-410bp) and 3) *C. caretta* R2SCOs size-selected at the same range (390-410bp). Stacks was used to obtain SNPs using the reference mode. Results showed that using the whole genome as a reference generated a large number of variant sites, but at low average coverage (15.1x ± 6.0). After running the populations module of Stacks, the R2SCOs generated more SNPs (14,610 versus 13,882) with a higher average coverage (Table S8).

Although all five backcrossed individuals could be detected using all three datasets, the genome-based analysis generated a significantly higher level of false-homozygotes for *E. imbricata*. In the backcrossed individuals against *C. caretta*, homozygotes for *E. imbricata* alleles are not expected and can be used as benchmark for detecting false homozygotes. When comparing the size-selected draft genome and the R2SCOs, the latter showed a lower rate of false-homozygotes, although not at a significant level (Fig. S13).

### New World Monkeys

ddRAD (2×150 bp) paired-end reads from *L. lagotricha, S. leucopus* and *S. flavius* (Valencia et al., 2018) were merged and analyzed using the R2SCO pipeline as described in the Methods. We selected a range of 205-230 bp for all three species, based on the number of reads divided by the number of expected loci per fragment size obtained from the digestion of a *Alouatta palliata* draft genome (Genbank ID GCA_004027835.1). More than 96% of the reads passed the preprocessing filtering workflow for each individual, but only between 21.4% and 40.6% were present within the selected size range (Table S9, Fig. S14). The number of accepted R2SCOs for *L. lagotricha, S. leucopus* and *S. flavius* were 23,291, 23,351 and 24,590, respectively (Table S10). The heterozygosity levels were similar in both individuals from each species (Table S11), and *L. lagotricha* was the most polymorphic species, followed by *S. flavius* and *S. leucopus*.

## 4. DISCUSSION

### A new strategy to build ddRAD-like locus catalogs

Here we present a carefully designed (R2SCO) pipeline, which proved to be a powerful tool to build reliable genome reduced-representation references for intra or interspecific analyses. Our study model involved five different species of sea turtles, but reference catalogs were built for each species independently. As the first step, in order to build high-quality locus catalogs that maximize the identification of variation for future population or other types of studies, individuals should ideally represent most of the diversity across the set of populations to be analyzed (Davey & Blaxter, 2010). Therefore, we selected two individuals for each species coming from the most genetically distant populations we could identify in our collection. For future studies, three or even more individuals from the same species can be used to build a reference, depending mostly on the tradeoff between costs (Table S12) and level of divergence expected.

The biggest novelty in our approach, in terms of methodology, is the use of overlapping 300 bp paired-end reads. Illumina sequences have very low error rates (Pfeiffer et al., 2018), however, base quality usually drops towards the end of the read. This is especially true for the longest Illumina reads (i.e. 300 bp paired-end reads from MiSeq), as also observed in our run quality controls (data not shown). This problem was completely solved by merging the reads, and quality was generally very high across the entire extent of the merged sequences. This was also verified in our quality filtering step, which removed very few sequences. The pattern of low removal rates remained the same when we increased the average score threshold to Q30.

With high-quality sequences representing the original fragment length, we could use the distribution of fragment lengths to evaluate the size selection performed in the lab and to perform a new size selection *in silico*, in order to homogenize coverages across loci and individuals. Our analysis of the fragment length distribution revealed huge variations across ddRAD libraries. However, even if in some cases we had to remove more than 80% of the sequences, we managed to select a range of well covered (>20x) loci across all five species, representing roughly 25,000 single-copy loci per individual. The R2SCO pipeline generated not only well-supported single-copy loci, but also gave indications of whether SNP calling would be able to retrieve all of the variation present in the alleles of each locus. The loci “removed” by our decision tree have issues of either biological (e.g. one allele out of range) or technical background (e.g. erroneous SNP calling, accumulation of sequencing errors, or alleles that are not mappable to representative locus sequences).

The consistency across individuals is also seen in the comparison of expected versus observed average locus coverage per length (Fig. S15). The putative alleles are compared to each other exhaustively (i.e. pairwise for all possible combinations), and a new script was designed to build clusters based on the pairwise alignments above the ancestral threshold. The exhaustive search for identity connections guarantees reproducibility of alleles clusterization within and between individuals. Since the data was significantly reduced to a set of high-quality and high-coverage sequences, the extended computational time demanded for exhaustive searches became a minor issue. We also performed the pairwise comparisons including putative alleles from all 10 individuals at the same time, and the alignment results were used for all intra-individual, intraspecific and interspecific analyses later on.

We also selected identity and coverage thresholds in an innovative way, using the identities generated by the pairwise comparisons to help establish ancestral as well as intraspecific thresholds. *T*_*intra*_ correlates negatively with the species heterozygosity, as it should be set to the initial point of the distribution of identities between orthologous alleles (Figure 3). In fact, *D. coriacea* and *L. olivacea* presented the highest *T*_*intra*_ and the lowest heterozygosity, while *C. mydas*, the most polymorphic species, presented the lowest *T*_*intra*_. *T*_*intra*_ is comparable to the value *M* in Stacks (Catchen et al., 2013), as it is set to avoid over-splitting of alleles, but should still keep the number of allowed differences low considering an intra-specific analysis. In contrast, *T*_*ances*_ is set to accommodate recent paralogy identity within or across species. The combination of both thresholds allowed us to keep clusters representing paralogous loci in our catalog and annotate them as putative paralogs within species. This way, we still perform comparisons of loci within or between species using lower thresholds (*T*_*ances*_), even if using an intra-specific identity threshold (*T*_*intra*_) for locus orthology classification for each species independently. For future analyses using the R2SCO rationale, we recommend *T*_*intra*_ to be always estimated for any given species, while *T*_*ances*_ can be kept at a very conservative value (e.g. 90%). We followed the latter recommendations for the new world monkey analysis as an example.

Given our analysis and comparisons to the genome, we believe that our set of R2SCOs for each species is better suited for future population studies than the draft genome available, when using ddRAD-like datasets. Although most of the loci were classified as single copy in both *C. mydas* R2SCOs and in the genome, we have identified large numbers of problematic loci that will most likely cause issues during genotyping of population data. We identified a variety of possible explanations, such as: presence of a second allele out of the size selected range, mostly due to internal indels; too many erroneous sequences within a locus; errors in homopolymeric or microsatellite regions; difficulties for the SNP caller to properly identify some longer or complex indel regions; issues with mapping for part of the reads that originally matched one allele within a locus; among others. Finally, even though the genome presents very high homology with both Cm individuals (>98%, Table S2), when simulating the ddRAD digestion *in-silico*, several loci were missing in the genome in comparison to Cm1 and Cm2. This is caused by mutations at the restriction sites, which are expected to happen more often between distant populations. In fact, the genome sample comes from an individual from the Indo-Pacific (Wang et al., 2013), in contrast to our two individuals from southwestern Atlantic populations.

### Sea turtles genetic variability

We used ∼5,5K putative single copy loci distributed across the sea turtle genomes to estimate heterozygosity levels per bp. Since our approach used entire sequences from merged paired-end reads, we could also use the invariant positions to estimate levels of genetic variation. The heterozygosity levels (0.028% and 0.031%) found for the two *C. mydas* individuals are slightly higher but not significantly different from the heterozygosity (AVG=0.024%, SD=0.018%) estimated across the draft *C. mydas* genome (Fitak & Johnsen, 2018). *Chelonia mydas* was substantially more variable compared to the other four sea turtle species, but heterozygosity levels can still be considered low for vertebrates. For example, in an analysis of three crocodilian genomes, Green et al. (2014) classified their heterozygosity levels (between 0.01% and 0.036%) as low when compared to avian and mammalian genomes. The least variable species, the American alligator *Alligator mississippiensis* (Daudin, 1802), presented genetic diversity values comparable to *E. imbricata*, the second most diverse species in our study (Figure 11). The very low genetic diversity observed for *L. olivacea* and *D. coriacea* compared to other sea turtle species is in accordance with their lower haplotypic diversity based on mitochondrial control region data for worldwide populations (Reid, Naro‐Maciel, Hahn, FitzSimmons, & Gehara, 2019). Another study using reduced-representation methods on six sea turtle species (Komoroske, Miller, & O’Rourke, 2019) also found *C. mydas* as the most variable and *D. coriacea* as the least variable species. The latter study -mostly based on Pacific populations -found *L. olivacea* as the second most variable sea turtle species, in contrast to our analysis of southwestern Atlantic populations. However, the authors point out to the fact that their largely different sample sizes might affect genetic variability estimation. Despite the use of reduced representation data, the heterozygosity values between studies are not directly comparable since Komoroske et al. (2019) has based their genetic diversity values on polymorphic positions only.

### Possible adjustments for future libraries and analysis limitations

By using the R2SCO approach, we produced a highly informative dataset that showed us the whole extension of fragment size distributions and differences across libraries and individuals. We were therefore able to detect several different issues in our libraries that could be evaluated and will be optimized in future experiments. For example, in spite of the use of high-precision equipment for size selection (i.e. BluePippin), our data revealed significant carryover of small fragments, as also seen in other studies (e.g. DaCosta and Sorenson, 2014). We believe that this is mostly due to the use of the MseI enzyme, a 4-bp cutter (recognition site: AATT) that cuts the genome in a very high frequency, generating large amounts of small fragments (Fig. S5). Small fragments were likely over amplified during the indexing PCR (Dabney & Meyer, 2012), which was performed before the size selection step. We hypothesize that the extreme levels of small fragments in the libraries might interfere in the BluePippin ability to remove them all. One simple future adjustment to avoid major carryover of small fragments is performing PCR indexing after the size selection step, like proposed in the original protocol (Peterson et al., 2012). Like in other studies, we changed the order of steps in an attempt to increase the reproducibility of the size selection step, avoiding posterior steps. Another sensible change involves using enzymes that do not produce such a biased ratio towards small fragments. Similarly to DaCosta and Sorenson (2014), the fragment analyzer analysis did not show any obvious presence of small fragments after our libraries were ready for sequencing (data not shown). Regardless of the library efficiency preparation, we believe that R2SCO datasets are powerful enough to allow for a very efficient *in-silico* adjustments of the ddRAD-like libraries, increasing the confiability of genotyping in any case where enough coverage is reached for a given size range across individuals.

Our approach presents a few limitations that should not be overlooked but can be mitigated. For example, paralogs that are very closely related and present up to two alleles in total are hard to be detected with just a few individuals. If alleles of such paralogs have identities above *T*_*intra*_ and coverage does not reach outlier levels, the only way to identify them is by using either a high-quality assembled genome or population data, since in the latter the proportion of heterozygous individuals should be greater for duplicated loci in comparison to single-copy loci at any given allele frequency (McKinney, Waples, Seeb, & Seeb, 2017). One other potentially important limitation of the method used is the lack of possibility to detect PCR duplicates. This can however be easily changed by adding methods that identify PCR duplicates through for example random nucleotides added in one of the Illumina indexing adaptors (as described in Hoffberg, Kieran, & Catchen, 2016). We took actions to minimize PCR duplicates, such as keeping PCR cycles to a maximum of 10 and starting each library with high amounts of DNA (1 µg), but we still cannot assess potential issues caused by technical duplicates generated doing amplification.

### Building R2SCOs for other species

Using sea turtle data, Chow et al. (2019) showed that reference-based SNP discovery using genomes of more closely related species allows for the identification of more SNPs. Similar to several other studies, they demonstrate the importance of establishing a proper reference for the target species. Downstream analysis has also been shown to be affected by the use of more distant reference genomes, causing reduced estimates of differentiation (Bohling, 2020). Galla et al. (2018) has demonstrated that for some types of analysis, a high-quality genome from a confamilial species can generate similar results that are relevant for conservation purposes (e.g. relatedness, nucleotide diversity and individual heterozygosity). Here we present a method to build a locus catalog that is comparable to results from a conspecific genome yet better suited for related species and even populations divergent from the genome. Our approach can potentially be transferred to any other species, as demonstrated here for previously published data from new world monkeys (Valencia et al., 2018). We also successfully tested it in birds, mammals and freshwater turtles (manuscripts in preparation).

We developed a pipeline that includes several new scripts, all of which were made available in a github page (see Data Accessibility Statement), and that include the use of SQLite databases. However, we understand the difficulties that many biologists face when trying to adapt scripts to their studies and while we don’t offer an automated pipeline in this article, we have designed scripts that should easily reproduce the R2SCO rationale starting from a locus catalog produced by Stacks (*Stacks2R2SCOs* workflow). We demonstrated here how restricting the size range to well covered fragment sizes increases mean coverage and general levels of polymorphism per locus in the final reference catalog.

Low coverage results in a decreased ability to confidently detect heterozygotes, affecting the analysis involving individual genotypes (Barbanti et al., 2020; Chow et al., 2019). Our results strongly indicate that a lot of variation is lost when accepting genotypes of loci with coverage below 15 reads. Sequencing at low coverage (e.g. <20x in average per locus) should especially be avoided when building the set of R2SCOs for a species, as the evaluation of heterozygosity as well as the potential technical issues regarding some loci will be crucial to help planning larger scale analyses (that will frequently involve shorter sequence reads) and will therefore decrease issues related to genotyping at the population level.

As pointed out by McCartney‐Melstad et al. (2019) and Paris et al. (2017), no single default value for the clustering threshold should be expected to be accurate for all studies or species. Our approach can help define empirical thresholds for any new species analyzed, with no need for previous knowledge regarding that species’ genetic variability. Existing methods for defining thresholds (Paris et al., 2017; Rochette & Catchen, 2017) seem to be largely robust and in our dataset they generated very similar identity thresholds as the R2SCO approach for four out of the five species. However, even though our analysis clearly defined a threshold of 99.1% for *D. coriacea*, the Stacks-based parameter testing remained inconclusive (Fig. S16).

Producing R2SCOs has one single requirement: sequencing overlapping paired-end reads. The analysis of the merged reads will allow researchers to properly evaluate the efficiency of library preparation (i.e. accuracy levels of digestion and ligation) as well as to select a size range with optimal coverage across individuals. Although a good understanding of the library outcome is essential to ensure genotype accuracy in downstream analysis, researchers should be attentive to several common points during strategy design and library preparation that will lead to high levels of removed reads in the pre-processing pipeline. We have listed several of these issues in Supplementary Table S13 and included cons and pros to different steps.

Finally, we recommend that researchers use informed assumptions about population representativeness to select specimens to build a locus catalog for a given study, independently of a focus on the entire species or on a set of populations only. In this way, the new reference set would not only be representative of the set of populations analyzed, but could also be used for a preliminary analysis of genetic distances among different populations, even with one single individual per population.

### Definition of the R2SCO term

We have designed a *de novo* locus catalog pipeline that incorporates information about coverage from merged sequences based on fragment size and uses the entire sequence as a putative allele. Sequences within each individual are compared based on two identity thresholds derived from pairwise comparisons of putative alleles, and differences between putative orthologous alleles are evaluated by a decision tree through a comparison against results from SNP calling. At the end of this pipeline, loci with accepted genotypes are considered as the set of reduced-representation single-copy orthologs (or R2SCOs, pronounced “Artuscos”). The idea to form a set of R2SCOs was inspired by BUSCO (Benchmarking Universal Single-Copy Orthologs, Seppey, Manni, & Zdobnov, 2019), but instead of representing orthologous single-copy coding genes among species, R2SCOs represent a set of orthologous loci produced by a certain combination of enzymes and a specific size range for a given species or set of species. R2SCOs from this study, for example, are defined as R2SCO-MseI-EcoRI-384-448, in which the last two numbers represent the *in-silico* size selection range.

## Conclusions

Coupling stepwise quality control and overlapping reads, this study presents a new approach to build ddRAD-like reference locus catalogs that also offers a detailed view over the libraries produced. Our empirical tests showed potential issues in the ddRAD size selection step and subsequent bioinformatic analyses. The use of long haplotypes, for some or all samples, can substantially improve the quality of the set of reference loci, thus potentially eliminating a series of biases incorporated during analysis that can strongly affect downstream analyses.

There is an ever-growing number of studies trying to overcome issues produced by RAD experiments and data (e.g. Barbanti et al., 2020; Chow et al., 2019; LaCava et al., 2020; Paris et al., 2017; Shafer et al., 2017). However, the issues caused by the technique are highly variable and seem to be specific to each experiment (DaCosta & Sorenson, 2014). The approach developed here proposes to use overlapping reads to gain power over the data created, helping scientists not only to produce high-quality reference sets of reduced-representation single-copy orthologs (R2SCOs), but also to evaluate each library created for a given species or population.

## Acknowledgements

Part of this work was supported by Alexander von Humboldt Foundation, Bonn, Germany, through a research grant to Sibelle T. Vilaca. This work was also supported financially by the Leibniz Institute for Zoo and Wildlife Research and the ANTIDOT project (Pépinière Interdisciplinaire Guyane, Mission pour l’Interdisciplinarité, CNRS). All sequencing and computer infrastructure utilized was part of the BeGenDiv Consortium. We are grateful to the Brazilian sea turtle conservation program Projeto TAMAR for collecting and providing part of the samples. We are similarly grateful to Prof. Fabricio Santos, who provided some of the Brazilian samples. We also thank Jilda Caccavo for a thorough English review of the text.

## Data Accessibility Statement

The codes in python are available through github at https://github.com/BeGenDiv/Driller_et_al_2021. The *Stacks2R2SCOs* workflow is available through https://git.imp.fu-berlin.de/begendiv/stacks2r2scos. The ddRAD data are available in the Sequence Read Archive through bioproject PRJNA######. The sets of R2SCOs representative sequences for each individual and comparison levels (species, Carettini, Cheloniidae, Chelonioidea) are available at Dryad doi:##.####/dryad.#####

## Author Contributions

C.J.M. and S.T.V. designed the study. B.T. and D.C. provided samples and financial support. S.T.V. performed the lab work. S.M. performed the sequencing. C.J.M, M.D. and L.S.A. analyzed the data. M.D. developed the accompanying scripts and produced most of the figures. T.C.V. supported the development of the decision tree. F.H. developed the code for clustering. C.J.M., M.D and L.S.A. wrote the manuscript with input from all co-authors.

## Supporting Information

## Supplementary Methods

## Supplementary Results

**Table S1:** Chimera detection results

**Table S2:** Levels of putative homology between 5 sea turtle species and the *Chelonia mydas* genome.

**Table S3:** Decision tree classifications of R2SCOs for each sample.

**Table S4:** Comparison of overlap between Stacks final catalog and R2SCOs locus classifications for each species.

**Table S5:** Number of missing loci between genome and *C. mydas* individuals

**Table S6:** Heterozygosity per bp, % of heterozygous loci and numbers of Chelonioidea-R2SCOs.

**Table S7:** Pairwise genetic distance (in percentage) matrix among 10 individuals.

**Table S8:** Stacks analysis using sea turtle data from Arantes et al. 2020.

**Table S9:** Sequence counts for each step of the preprocessing workflow for Valencia et al. data.

**Table S10:** Number of loci obtained by the R2SCO pipeline for three new world monkey species.

**Table S11:** Heterozygosity per bp, % of heterozygous loci and numbers of R2SCOs per individual, including only accepted loci with coverage above 15 for Valencia et al. data.

**Table S12:** Estimated costs for producing high-quality R2SCOs for a single individual.

**Table S13:** Pros and Cons of alternative methods for building ddRAD-Seq libraries.

**Figure S1**. Fragment size distribution of merged and preprocessed reads for 10 sea turtle individuals from 5 species across four sequencing runs.

**Figure S2:** Read preprocessing workflow.

**Figure S3**: Decision tree for classification of loci clustered within individuals using the R2SCO pipeline

**Figure S4:** Workflow for chimera detection.

**Figure S5:** *C. mydas* genome *in silico* digestion including the *in-silico* size range selected

**Figure S6:** Stacks2R2SCOs workflow to filter the stacks catalog to generate an R2SCO like filtered catalog.

**Figure S7:** Results of preprocessing analysis within the selected size range (384-448 bp).

**Figure S8:** Megan trees from preliminary contamination analysis.

**Figure S9:** Coverage distribution across loci within each individual.

**Figure S10:** Number of loci, total coverage per fragment size and average coverage per locus across the selected size range for each individual.

**Figure S11:** Intra-individual heterozygosity rates across different phylogenetic levels of R2SCOs.

**Figure S12:** Heatmap of pairwise genetic distances across all individuals from the five sea turtle species.

**Figure S13:** Ratio of incorrectly assigned genotypes using different references (*C. mydas* genome without treatment, *C. mydas* genome size selected and R2SCOs for *C. caretta* size selected) for backcrossed hybrids (*C. caretta* x *E. imbricata* hybrid) X C. caretta analyzed in Arantes et al., 2020b. R2SCO reference showed the lowest rate of false-homozygotes for E. imbricata (homozygotes for *E. imbricata* alleles are not expected in the backcrossed individuals) compared to the other references, although not at a significant level.

**Figure S14:** Heterozygosity levels for R2SCOs from primates from Valencia et al. 2018.

**Figure S15:** Boxplots representing the distribution of coverage per locus across fragment size.

**Figure S16**. Difference in number of new polymorphic loci added for each iteration of *M* for *Dermochelys coriacea*.

**Figure S17:** Circos plots of comparisons of two *C. mydas* individuals (Cm1 and Cm2) and their combined loci (Cm species) against the genome clusters of digested sequences.

## Notes

https://github.com/BeGenDiv/Driller_et_al_2021

https://git.imp.fu-berlin.de/begendiv/stacks2r2scos

### Competing Interest Statement

The authors have declared no competing interest.

### Summary of Updates

The new version focuses almost entirely on the R2SCO method design, description, implementation and interpretation, where sea turtles are used almost exclusively as a model to demonstrate the method. We have therefore changed the title of the manuscript accordingly. Three major changes can be found: 1) addition of detailed comparisons of the R2SCO pipeline against Stacks; 2) development of a workflow to help converting Stacks-generated locus catalogs into sets of R2SCOs; 3) use of different datasets to show the usage (as reference) and flexibility (regarding other vertebrate species) of the R2SCOs pipeline.

