## Supporting_Info for "Achieving high-quality ddRAD-like reference catalogs for non-model species: the power of overlapping paired-end reads"

Contents

Supplementary Methods……………...……………..…….……….………………………………...2
Supplementary Results..……………...……………..…….……….………………….....…………..5
Supplementary Tables..…………………………………………………………....…………………6
Supplementary Figures..……………....…………………..……………………………………….14

### Supplementary Methods

### Chimera detection workflow

First, fragments originated from each digested sequence were independently mapped to the *C. mydas* reference genome and checked for continuity, e.g. if the fragments of one sequence could not be mapped in the expected order (same orientation and 100 bp proximity) within the same scaffold the sequence was considered to be potentially chimeric. In order to avoid inflating the chimera estimations due to incorrect mappings of small fragments, the potential chimeras were mapped again with bowtie2 using the original (non-digested) sequence. Only sequences which did not align end-to-end or did not map at all were classified as chimeras, otherwise they were considered as incomplete digestion. The workflow for chimera detection can be found in Fig. S4.

### R2SCO pipeline details

#### *Thresholds definition*

*Minimum count per putative allele* (*m*)

The value of m represents the minimum coverage per unique sequence within an individual that should represent a putative allele. This value should be as low as possible, assuming that artifactual sequences rarely reach *m*. Our approach to choose *m* was based on the distribution of identical sequences with coverages 1-20 (data not shown). As our samples mostly did not pass average coverage of 50 reads per locus, the frequency of artifacts reaching m=3 was rather low and easily identifiable (see Fig. 5 for the analysis of replicate runs).

*Minimum locus coverage* ($COV_{min}$)

The minimum coverage per locus within an individual helps selecting loci within individuals with enough coverage to yield reliable genotypes. The chosen value of $COV_{min}$ was based on empirical analysis of replicates (Fig. 5). It is not used for defining R2SCOs since a locus can be defined by a single allele. Empirical tests in our dataset have shown that the great majority of heterozygous loci could be identified as such in both replicates when these have reached a minimum coverage of 15.

*Maximum locus coverage* ($COV_{max}$)

The maximum locus coverage is the limit between the distribution of sequence coverages of single-copy loci and outlier coverages that mostly represent paralogs. $COV_{max}$ per locus was chosen for each individual independently. This number was based on the upper end of the coverage distribution per locus, after clustering putative alleles based on $T_{ances}$. Bars (as in Fig. S9) are coloured according to the number of alleles present in the cluster (1 allele, 2 alleles, Paralog Individual - when more than 2 putative alleles are present, and Paralog Tintra - when the identity between 2 alleles is below $T_{intra}$). The threshold line is chosen manually at the upper end of the distribution of 1-2 alleles in order to mostly identify loci that have coverage at levels of putative paralogs - i.e. outliers. We have performed several tests, using likelihood, binomial distribution, and others, and none of them seem to produce reasonable results across all individuals, excluding in some cases large amounts of loci with 1 or 2 alleles with no indication of paralogy.

*Ancestral identity threshold* ($T_{ances}$)

The ancestral threshold represents a minimum identity that clusters recent paralogous loci. It can be based on the distance between the most distant samples or in case of an intra-specific analysis, it can be a conservative value that would avoid the successful mapping of sequences to a closely-related paralogous locus (e.g. 90%). In our case, in which we analyzed 5 species with increasing phylogenetic distances, it was reasonable to use the distribution of distances between each of the 4 other species and *D. coriacea* as a conservative way to identify paralogy that happened after the first split of sea turtles (as shown in Fig. 3). For intraspecific or closely-related species analysis, our recommendation lies on avoiding cross-matching of paralogy via mapping. This means that, in case a locus is covered by one individual, and it is an orphan of an orthologous locus in the reference, the mapping might happen to the closest paralogous locus. The ancestral threshold should be such as to avoid that these loci are accepted in the reference. Usually, 90% avoids cross mapping end-to-end, but one might want to test specific cases, in case there is an interest in increasing the ancestral threshold.

*Intra-specific identity threshold* ($T_{intra}$)

The identity threshold within species represents an identity value among sequences that clusters the great majority of orthologous alleles and excludes the great majority of paralogs. This threshold is obtained by comparing identities among putative alleles within and between individuals from the same species. For each species, the pairwise comparison between putative alleles within individuals (i.e. any two alleles from a heterozygous locus or any two alleles from paralogous loci with identity > $T_{ances}$) and between individuals (i.e. mostly orthologous alleles, in many cases identical, as long as both individuals covered the locus) helped establishing the intra-specific threshold ($T_{intra}$), above which most alleles should belong to orthologous loci (Figure 5).

***R2SCO pipeline steps iii-x***

The step *iii* is performed by the dereplicateFQ.py script and generates a fastq file containing only de-replicated (=unique) sequences for which one of the original IDs is randomly chosen among the identical sequences, followed by the total count of identical sequences. The phred quality scores are picked based on the highest score for each position after comparing the quality of all identical sequences. For defining the putative alleles in step *iv*, a minimum count per de-replicated sequence is chosen (analogous to parameter *m* from the program Stacks) that should represent each putative allele in the selected range. Sequences with a frequency less than the chosen threshold are ignored, since they should represent either sequencing errors or very low covered loci. The low-frequency sequences will be used at a later step for mapping to the putative loci, similarly to the *secondary reads* of Stacks. For step *v*, putative alleles from all individuals were concatenated into one file and aligned pairwise using vsearch --allpairs_global (v2.8.6) with an identity defined by $T_{ances}$ (here 0.90). Additionally, the parameter --iddef 3 was set, which treats each extended gap as a single difference. Finally, in step *vi* clusters are formed using the script clusterFromPairs.py for single-linkage clustering and the $T_{ances}$ identity value. Each cluster represents a homologous ddRAD locus (note that homology here includes both orthology and paralogy) within the genome. Clusters were generated within each sample, within each species and across multiple species.

In step *vii*, instead of mapping merged and pre-processed sequences to a consensus sequences, we have mapped the sequences to a multi-fasta file containing all putative allele sequences for a given individual using bowtie2 (parameters: --mp 10 --rdg 9,1 --rfg 9,1 --score-min L,-1,-0.2). For this step, the merged sequences were allowed to be 5% shorter or 5% longer than our size range limits in order to identify the presence of uncalled alleles with small indels. Subsequently, reads were considered to be part of a locus if they mapped to any of the putative alleles belonging to that locus. In order to perform a SNP calling in step *viii*, we gathered all reads of a certain locus and re-mapped them to either the longest (if alleles presented different lengths) or the most frequent putative allele of a given locus, using the same parameters as before. Variable positions were identified using the mpileup module of bcftools v.1.9 (Narasimhan et al., 2016) followed by the call module of bcftools with the additional parameters -mv -

Ob. In step *ix*, the VCF file containing both reference and alternative bases was used to reconstruct haplotypes using the script buildHaplotypes.py. All information obtained so far was then used for locus classification through the decision tree.

**Decision tree**

The input of the decision tree consists of each locus for a given individual as defined by the clustering of putative alleles based on pairwise comparisons (step *vii*). If any putative allele is successfully blasted (blastn with e-value < 10e-50 against nt) against a non-chordata sequence, the correspondent locus is considered to belong to some sort of contamination and is annotated as "Non-turtle". Putative endogenous loci are then tested for potential paralogy and can be classified as three different types of paralogs: 1) Paralog Individual, if both numbers of putative alleles and reconstructed haplotypes are higher than two, or if the number of putative alleles is above four (in which case SNP calling could fail as it largely violates the assumption of diploidy); 2) Paralog Coverage, if the sum of read counts mapped to each putative allele belonging to a locus is above the COVmax threshold for a given individual and 3) Paralog $T_{intra}$, if the pairwise comparison between any two alleles of a given locus yielded an identity below the $T_{intra}$ threshold established for the species. As classifications are performed stepwise, each locus can only obtain one single classification, after which the decision tree will re-start by evaluating the next locus.

Once annotation steps of non-Turtle and Paralogs are finalized, the next steps of the decision tree evaluate the reliability of loci based on two aspects: 1) The quality of reads re-mapping to the locus sequence representative and 2) The comparison between the putative alleles obtained from the merged sequences de-replication (step *vi*) and the reconstructed haplotypes obtained after SNP calling (step *ix*). There are five locus removal steps, which can be briefly explained in the following ways: Removed 1) If a significant amount of sequences (>10% or >2) do not (re-)map from the start to the end of the locus (i.e. from restriction site to restriction site); Removed 2) If the number of SNPs identified between putative alleles is different from the number of SNPs identified by SNP calling; Removed 3) If there are reconstructed haplotypes out of the size range defined *in silico* (which would not be present among the putative alleles due to step *ii*); Removed 4) If the number of haplotypes is higher than the number of alleles; Removed 5) If the number of alleles is higher than the number of haplotypes and frequencies were still high enough to reject a persistent sequencing error among alleles.

In cases in which more alleles than haplotypes are found but counts were very low for the extra alleles, the haplotype genotype was accepted (Accepted Genotype from Haplotype) assuming that putative alleles were formed by persistent errors that reached the minimum frequency *m*. In all other cases, the genotype from alleles - which should be identical to the one from haplotypes - was accepted (Accepted Genotype from Allele). In those cases in which the loci with accepted genotypes had low frequency (i.e. $<COV_{min}$), the locus was accepted but considered as low frequency (Accepted Genotype low frequency). Only loci with accepted genotypes (including low frequency ones) were included in the set of R2SCOs from an individual. A visualisation of the decision tree can be seen in Fig. S3.

**Further considerations about R2SCOs**

R2SCOs can represent different levels of comparisons: set of populations, whole species, set of species. The set of R2SCOs coming from the same enzyme combination and size range may change across these levels, as mutations at the restriction sites may cause entire locus dropouts and some loci may be annotated as paralogs in different species or populations. Therefore, an R2SCO set within or between species is also a locus catalog under construction and must be re-evaluated after each new experiment.

***In-silico* size selection**

The awk command used to select sequences between 348 and 448 bp:

```
awk 'BEGIN {OFS = "\n"} {header = $0 ; getline seq ; getline qheader ; getline qseq ; if (length(seq) >= 384 && length(seq) <= 448) {print header, seq, qheader, qseq}}' < input.fastq  > output.fastq;
```

### Supplementary Results

#### Reference Genome x R2SCOs

There was a non-neglectable level of missing loci (between 1,212 and 4,229 loci) in both the genome and the individuals analyzed (Table S5). When using Cm1 - the individual with the poorest coverage - as a parameter for locus presence, the numbers of missing loci was much higher in the genome (3,031) compared to Cm2 (1,212), indicating that the latter has a much higher level of locus overlapping with Cm1.

There is also a high proportion of removed loci for Cm1 and Cm2 (11.28% and 7.23%, respectively), most of which were found as singletons in the genome. Those loci are mostly variable but with some issue on the genotyping step identified by the decision tree. Looking more closely at the possible issues, we identified a variety of possible explanations, such as: presence of a second allele out of the size selected range, mostly due to internal indels; too many erroneous sequences within a locus; errors in homopolymeric or microsatellite regions; difficulties for the SNP caller to properly identify some longer or complex indel regions; issues with mapping for part of the reads that originally matched one allele within a locus; among others.

As seen in Figure 7B, paralogy is very minor among the loci within the selected size range (1.04% of the genome digested fragments). However, looking more closely at the putative paralogs and their relation to the genome (Fig. S17), a substantial part of the paralogs defined by the $T_{intra}$ threshold are present in a single copy in the genome. The latter amounts for roughly 2,62% for Cm1 and 2,56% for Cm2 of loci that appear as single-copy in the genome. These values are probably slightly overestimated because they exclude all the removed loci from Cm. Nevertheless, the values approach our goal of leaving a tail of about 2% of all the orthologous loci with alleles identity falling under $T_{intra}$, when defining this threshold. In contrast, loci with three or more alleles identified after the decision tree (Paralog individual) seem to be present in the genome in the following decreasing order of frequency: missing, paralogs and singleton.

While most of the loci present in three or more copies (P3+) in the genome also showed up as paralogs in the Cm individuals, the majority of the genome loci with a second similar locus (P2) in the assembly were accepted as single copy in the two individuals (Figure 7B). One possible explanation is a small level of haplotype retention in the genome assembly, common to draft genomes.

### Supplementary tables

**Table S1.** Chimera detection results

| sID | merged | totalDigested | noInternalRS | unmapped | incompDig | chimeric |
|---|---|---|---|---|---|---|
| Cc1 | 3,439,549 | 202,185 | 3,237,364 | 21,531 | 142,132 | 38,522 |
| Cc2 | 3,636,115 | 415,697 | 3,220,418 | 47,536 | 161,110 | 207,051 |
| Dc1 | 3,275,771 | 356,462 | 2,919,309 | 87,643 | 139,123 | 129,696 |
| Dc2 | 2,606,818 | 403,976 | 2,202,842 | 98,230 | 104,997 | 200,749 |
| Ei1 | 4,731,081 | 444,239 | 4,286,842 | 64,126 | 166,055 | 214,058 |
| Ei2 | 2,332,021 | 350,998 | 1,981,023 | 31,215 | 304,777 | 15,006 |
| Lo1 | 1,725,492 | 313,020 | 1,412,472 | 29,655 | 139,462 | 143,903 |
| Lo2 | 1,568,494 | 193,890 | 1,374,604 | 27,573 | 49,258 | 117,059 |
| Cm1 | 2,848,253 | 1,051,900 | 1,796,353 | 27,197 | 976,316 | 48,387 |
| Cm2 | 3,933,845 | 352,632 | 3,581,213 | 16,833 | 237,197 | 98,602 |

**Table S2.** Levels of putative (cumulative) homology between 5 sea turtle species and the *Chelonia mydas* genome with an additional blast against *Chrysemys picta*.

| Sample | #reads | Mapped CheMyd | Blasted CheMyd | Blasted ChrPic |
|---|---|---|---|---|
| **Cc1** | 50,000 | 97.82% | 98.46% | 99.27% |
| **Cc2** | 50,000 | 97.77% | 98.39% | 99.06% |
| **Lo1** | 50,000 | 98.11% | 98.77% | 99.51% |
| **Lo2** | 50,000 | 97.13% | 97.94% | 98.66% |
| **Ei1** | 50,000 | 96.33% | 96.85% | 97.74% |
| **Ei2** | 50,000 | 97.23% | 97.96% | 98.68% |
| **Cm1** | 50,000 | 98.20% | 98.79% | 99.39% |
| **Cm2** | 50,000 | 98.21% | 98.77% | 99.54% |
| **Dc1** | 50,000 | 94.21% | 95.58% | 97.12% |
| **Dc2** | 50,000 | 95.50% | 97.31% | 98.72% |

Note: Cumulative matching percentages are shown after each step.

**Table S3.** Decision tree classifications of R2SCOs for each sample.

| Sample | Total | Total Accepted | Total Paralogs | Total Removed | Accepted Allele | Accepted Haplotype | Accepted Low freq | Non-Turtle | Paralog Individual | Paralog Coverage | Paralog Tintra | Removed 1 | Removed 2 | Removed 3 | Removed 4 | Removed 5 |
|---|---|---|---|---|---|---|---|---|---|---|---|---|---|---|---|---|
| Cc1 | 25,904 | 24,563 | 462 | 854 | 23,532 | 141 | 890 | 25 | 163 | 89 | 210 | 104 | 250 | 130 | 199 | 171 |
| Cc2 | 25,107 | 23,683 | 360 | 1,058 | 19,859 | 89 | 3,735 | 6 | 152 | 46 | 162 | 93 | 332 | 108 | 390 | 135 |
| Lo1 | 23,966 | 22,975 | 288 | 702 | 16,540 | 34 | 6,401 | 1 | 129 | 31 | 128 | 54 | 184 | 52 | 330 | 82 |
| Lo2 | 21,805 | 20,725 | 254 | 620 | 12,991 | 269 | 7,465 | 206 | 125 | 22 | 107 | 29 | 251 | 23 | 209 | 108 |
| Ei1 | 24,678 | 22,600 | 431 | 1,634 | 15,951 | 57 | 6,592 | 13 | 215 | 29 | 187 | 102 | 628 | 125 | 691 | 88 |
| Ei2 | 25,505 | 23,278 | 462 | 1,716 | 17,042 | 130 | 6,106 | 49 | 211 | 46 | 205 | 85 | 693 | 125 | 686 | 127 |
| Cm1 | 25,378 | 21,856 | 638 | 2,861 | 17,943 | 24 | 3,889 | 23 | 274 | 23 | 341 | 227 | 914 | 311 | 1,286 | 123 |
| Cm2 | 27,054 | 24,329 | 752 | 1,956 | 23,382 | 85 | 862 | 17 | 290 | 76 | 386 | 255 | 662 | 385 | 470 | 184 |
| Dc1 | 25,095 | 24,197 | 239 | 548 | 20,773 | 34 | 3,390 | 111 | 107 | 22 | 110 | 44 | 105 | 64 | 237 | 98 |
| Dc2 | 25,346 | 24,543 | 222 | 581 | 22,220 | 83 | 2,240 | 0 | 84 | 36 | 102 | 39 | 101 | 75 | 215 | 151 |

**Table S4. Comparison of overlap between Stacks final catalog and R2SCOs locus classifications for each species (overlaps are shown in %)**

| | Cc | Lo | Ei | Cm | Dc |
|---|---|---|---|---|---|
| **stacks_r2scos** | 93.20 | 94.32 | 91.61 | 89.75 | 93.60 |
| **stacks_r2scosAcceptedLF** | 87.78 | 90.44 | 82.01 | 77.84 | 90.74 |
| **stacks_r2scosRemoved** | 5.42 | 3.88 | 9.60 | 11.91 | 2.86 |
| **stacksOnly** | 1.10 | 0.78 | 1.49 | 01.07 | 0.88 |
| **r2scosOnly** | 5.69 | 4.90 | 6.91 | 9.18 | 5.52 |

**Table S5. Number of missing loci between genome and *C. mydas* individuals**

| | *Missing in* | | |
|---|---|---|---|
| *From* | Genome | Cm1 | Cm2 |
| Genome | - | 4,229 | 2,955 |
| Cm1 | 3,031 | - | 1,212 |
| Cm2 | 3,447 | 2,900 | - |

**Table S6.** Heterozygosity per bp, % of heterozygous loci and numbers of Chelonioidea-R2SCOs used per individual, including only accepted loci with coverage above 15.

| Species | Sample | % Heterozygosity /bp | SNPs per Locus | % Heterozygous Loci | # Heterozygous Loci | # Loci | # Polymorphic Sites | Summed Length |
|---|---|---|---|---|---|---|---|---|
| **Cc** | **Cc1** | 0.075 | 0.312 | 23.09 | 475 | 2,057 | 641 | 859,492 |
| **Cc** | **Cc2** | 0.077 | 0.320 | 23.22 | 477 | 2,054 | 658 | 858,105 |
| **Cc** | **Cc1-Cc2** | 0.152 | 0.637 | 40.38 | 821 | 2,033 | 1,295 | 849,549 |
| **Lo** | **Lo1** | 0.033 | 0.139 | 11.86 | 245 | 2,065 | 287 | 862,990 |
| **Lo** | **Lo2** | 0.037 | 0.158 | 12.91 | 214 | 1,657 | 261 | 703,338 |
| **Lo** | **Lo1-Lo2** | 0.036 | 0.155 | 12.50 | 206 | 1,648 | 255 | 699,514 |
| **Ei** | **Ei1** | 0.109 | 0.456 | 30.99 | 639 | 2,062 | 940 | 861,746 |
| **Ei** | **Ei2** | 0.101 | 0.421 | 29.07 | 596 | 2,050 | 863 | 856,786 |
| **Ei** | **Ei1-Ei2** | 0.108 | 0.451 | 30.26 | 616 | 2,036 | 919 | 850,907 |
| **Cm** | **Cm1** | 0.291 | 1.219 | 54.31 | 1,108 | 2,040 | 2,486 | 852,990 |
| **Cm** | **Cm2** | 0.292 | 1.221 | 52.39 | 1,076 | 2,054 | 2,508 | 859,128 |
| **Cm** | **Cm1-Cm2** | 0.322 | 1.347 | 57.09 | 1,152 | 2,018 | 2,718 | 844,113 |
| **Dc** | **Dc1** | 0.027 | 0.112 | 9.47 | 196 | 2,069 | 232 | 862,932 |
| **Dc** | **Dc2** | 0.028 | 0.118 | 9.83 | 203 | 2,066 | 243 | 861,823 |
| **Dc** | **Dc1-Dc2** | 0.030 | 0.124 | 10.52 | 217 | 2,063 | 255 | 860,529 |

**Table S7.** Pairwise genetic distance (in percentage) matrix among 10 individuals from the five sea turtle species, based on total differences between the consensus sequences of shared loci averaged by length.

| Sample | Cc1 | Cc2 | Cm1 | Cm2 | Dc1 | Dc2 | Ei1 | Ei2 | Lo1 | Lo2 |
|---|---|---|---|---|---|---|---|---|---|---|
| **Cc1** | - | 0.140 | 2.292 | 2.293 | 4.662 | 4.667 | 1.064 | 1.063 | 0.945 | 0.947 |
| **Cc2** | 0.140 | - | 2.293 | 2.294 | 4.660 | 4.663 | 1.066 | 1.064 | 0.948 | 0.950 |
| **Cm1** | 2.292 | 2.293 | - | 0.297 | 4.427 | 4.429 | 2.313 | 2.314 | 2.352 | 2.365 |
| **Cm2** | 2.293 | 2.294 | 0.297 | - | 4.442 | 4.445 | 2.318 | 2.321 | 2.356 | 2.370 |
| **Dc1** | 4.662 | 4.660 | 4.427 | 4.442 | - | 0.027 | 4.676 | 4.677 | 4.700 | 4.718 |
| **Dc2** | 4.667 | 4.663 | 4.429 | 4.445 | 0.027 | - | 4.681 | 4.682 | 4.703 | 4.723 |
| **Ei1** | 1.064 | 1.066 | 2.313 | 2.318 | 4.676 | 4.681 | - | 0.100 | 1.129 | 1.136 |
| **Ei2** | 1.063 | 1.064 | 2.314 | 2.321 | 4.677 | 4.682 | 0.100 | - | 1.133 | 1.140 |
| **Lo1** | 0.945 | 0.948 | 2.352 | 2.356 | 4.700 | 4.703 | 1.129 | 1.133 | - | 0.027 |
| **Lo2** | 0.947 | 0.950 | 2.365 | 2.370 | 4.718 | 4.723 | 1.136 | 1.140 | 0.027 | - |

**Table S8.** Stacks analysis using sea turtle data from Arantes et al. 2020.

| **Stacks Analysis** | **Number of loci obtained in gstacks** | **Number of loci after populations filters (r=0.4, p=3)** | **Number of Variant Sites** | **Coverage (mean ± stdev)** |
|---|---|---|---|---|
| Reference *C. mydas* genome without treatment | 332,054 | 67,328 | 143,438 | 15.1 ± 6.0 |
| Reference *C. mydas* genome size selected (390-410 bp) (Arantes et al 2020) | 23,406 | 4,534 | 13,882 | 39.8 ± 22.2 |
| Reference R2SCOs from *C. caretta* size selected (390-410 bp) | 7,187 | 4,409 | 14,610 | 42.5 ± 24.1 |

**Table S9.** Sequence counts for each step of the preprocessing workflow for Valencia et al. data.

| **Species** | **Sample ID** | **Demultiplexing** | **Trimmed** | **Merged** | **Non-Digested+Digested** | **Q>=30*** |
|---|---|---|---|---|---|---|
| ***Lagothrix lagotricha*** | DIFI1514_ref | 4,874,470 | 4,873,372 | 4,753,930 | 4,755,472 | 4,685,544 (96%) |
| | LAGORILEY_ref | 4,484,167 | 4,482,573 | 4,447,725 | 4,456,965 | 4,419,038 (98%) |
| ***Saguinus leucopus*** | SL012_ref | 4,111,658 | 4,110,022 | 4,082,304 | 4,109,940 | 4,073,396 (99%) |
| | SL127_ref | 4,131,620 | 4,130,308 | 4,087,635 | 4,095,982 | 4,052,003 (98%) |
| ***Sapajus flavius*** | CPB429_ref | 3,311,659 | 3,309,973 | 3,278,136 | 3,284,629 | 3,250,562 (98%) |
| | CPB462_ref | 4,444,294 | 4,442,638 | 4,392,442 | 4,393,482 | 4,342,508 (98%) |

*Quality filter was performed after trimming 8bp from both ends. % Percentages refer to the Demultiplexed sequences.

**Table S10.** Number of loci obtained by the R2SCO pipeline for three new world monkey species.

| | *Saguinus leucopus* | | *Lagothrix lagotricha* | | *Sapajus flavius* | |
|---|---|---|---|---|---|---|
| **Classifications** | **SL127** | **SL012** | **LAGORILEY** | **DIFI1514** | **CPB429** | **CPB462** |
| Possible contamination | 0 | 0 | 0 | 0 | 0 | 0 |
| Paralog_Individual | 364 | 267 | 237 | 353 | 281 | 349 |
| Paralog_Coverage | 100 | 61 | 27 | 44 | 35 | 67 |
| Paralog_Tintra | 179 | 149 | 343 | 254 | 255 | 249 |
| Low_freq | 1,790 | 9,227 | 2,014 | 1,067 | 2,238 | 1,110 |
| Accepted_Genotype_Allele | 24,195 | 14,448 | 20,843 | 2,4667 | 21,392 | 2,5404 |
| Removed1 | 284 | 295 | 465 | 326 | 333 | 282 |
| Removed2 | 267 | 1001 | 664 | 323 | 687 | 312 |
| Removed3 | 245 | 146 | 270 | 327 | 218 | 287 |
| Removed4 | 355 | 558 | 1081 | 315 | 1050 | 392 |
| Removed5 | 385 | 243 | 643 | 430 | 487 | 420 |
| Accepted_Genotype_Haplotype | 2,830 | 1,427 | 3,502 | 3,069 | 3,603 | 2,746 |
| Total | 30,994 | 27,822 | 30,089 | 31,175 | 30,579 | 31,618 |
| % of accepted+low-freq/total | 92,97% | 90,22% | 87,60% | 92,39% | 89,06% | 92,54% |

**Table S11.** Heterozygosity per bp, % of heterozygous loci and numbers of R2SCOs per individual, including only accepted loci with coverage above 15 for Valencia et al. data.

| **Species** | **Sample** | **% Heterozy-gosity/bp** | **SNPs per Locus** | **% Heterozy-gous Loci** | **# Heterozy-gous Loci** | **# Loci** | **# Polymorp hic Sites** | **Summed Length bp** |
|---|---|---|---|---|---|---|---|---|
| ***Lagothrix lagotricha*** | DIFI1514 | 0.163 | 0.381 | 30.701 | 8,843 | 28,803 | 10,975 | 6,715,726 |
| | LAGORILEY | 0.173 | 0.403 | 32.998 | 8,698 | 26,359 | 10,626 | 6,147,975 |
| | DIFI1514-LAGORILEY | 0.181 | 0.422 | 31.093 | 7,242 | 23,291 | 9,830 | 5,428,328 |
| ***Saguinus leucopus*** | SL012 | 0.068 | 0.159 | 13.425 | 3,370 | 25,102 | 3,992 | 5,851,862 |
| | SL127 | 0.098 | 0.228 | 18.806 | 5,419 | 28,815 | 6,571 | 6,718,039 |
| | SL012-SL127 | 0.091 | 0.211 | 16.843 | 3,933 | 23,351 | 4,937 | 5,442,415 |
| ***Sapajus flavius*** | CPB429 | 0.135 | 0.313 | 24.962 | 6,798 | 27,233 | 8,540 | 6,354,678 |
| | CPB462 | 0.127 | 0.297 | 22.146 | 6,480 | 29,260 | 8,684 | 6,823,171 |
| | CPB429-CPB462 | 0.155 | 0.361 | 24.677 | 6,068 | 24,590 | 8,877 | 5,735,,413 |

**Table S12. Estimated costs for producing high-quality R2SCOs for a single individual.**

| | Specifications | Price | Price per sample |
|---|---|---|---|
| **Digestion** | | | |
| NEB 10x CutSmart Buffer | | € - | € - |
| EcoRI-HF (20 U/µl) | EcoRI HF 10.000 U - 10 U / sample | € 50,00 | € 0,05 |
| MseI (20 U/µl) | MseI 500 U - 10 U / sample | € 71,00 | € 1,42 |
| **Ligation** | | | |
| Adapter P1 (5 µM) | Oligos up (€ 27,62) and down (€ 30,56) - 0,05 µmol | € 58,18 | € 0,014 |
| Adapter P2 (5 µM) | Oligos up (€ 25,66) and down (€ 21,25) - 0,05 µmol | € 46,91 | € 0,011 |
| 10x Ligase Buffer | | € - | € - |
| T4 DNA ligase (400 U/µL) | T4 DNA ligase 20.000 U - 100 U / sample | € 68,00 | € 0,34 |
| **Indexing PCR** | | | |
| 5x Polymerase Buffer | | € 22,80 | € 0,57 |
| dNTP's | | € 11,65 | € 0,29 |
| Polymerase | | € 45,60 | € 1,14 |
| PCR Primer 1 (10 µM) | 0,05 µmol | € 31,54 | € 0,015 |
| PCR Primer 2 (10 µM) | 0,05 µmol | € 34,97 | € 0,017 |
| **Size selection** | | | |
| BluePippin cassette (1.5% agarose) | Each cassette contains 5 lanes | € 73,51 / cassette | € 14,7023 / lane |
| **Product purification** | | | |
| Magnetic beads | 100 ul beads / sample | € 8,45 / mL | € 0,84 |
| **Product verification** | | | |
| Agarose gel | | € 1,00 | € 1,00 |
| **Product quantification** | | | |
| Qubit | 2 DNA quantification / sample | € 1,12 | € 2,23 |
| TapeStation Automated Electrophoresis System | Agilent Technologies (Price per lane) | € 5,22 | € 5,22 |
| **Disposables** | | | |
| Tips, tubes | | € 5,00 | € 5,00 |
| **Sequencing** | | | |
| MiSeq v3 600 cycles (25 M reads) | 3 M 2x300 bp reads per sample for 30X coverage | € 1.792,82 | € 215,14 |
| | | **TOTAL** | € 233,31 |

**Table S13. Pros and Cons of alternative methods for building ddRAD-Seq libraries**

| | Methods | Pros | Cons |
|---|---|---|---|
| **Produce of a reference Catalog using R2SCOs** | R2SCOs catalog building described in this paper | Provide a reliable set of reference loci and preliminary information about the genetic diversity of the species. Allow testing crucial steps in the ddRAD protocol (as the combination of enzymes and size selection) for a small number of representative samples, which result in a more accurate planning of the final library containing the complete set of samples and, consequently, better quality and useful ddRAD dataset. | Extra time (up to a month) to prepare the R2SCOs independently of the large ddRAD-like libraries; further costs (~250 EUR per individual used in the reference) |
| **In-vitro Size Selection** | Blue Pippin, Pippin Prep or Other methods for precision size selection excision (e.g. eGel) | Allows very precise size selection (although it does not seem to be exactly reproducible and prone to small fragment retention in cases where library are dominated by small fragments) | More expensive than common gel extraction or beads purification |
| | Common agarose gel excision | Low cost and of common use in any molecular laboratory | Low precision and optimization needed to prevent the carryover of small fragments |
| | Beads purification | Low cost after optimized | Needs extensive optimization to reach a somewhat precise range; |
| **Sequencing** | MiSeq platform | Generates longer sequences (up to 600 bp) that yield overlapping reads for larger size ranges (up to 550bp); reference R2SCOs for 2-3 individuals per species can use half or the entire run (easier to fit into a run) | Higher costs per bp |
| | NextSeq, HiSeq or NovaSeq platforms | Lower sequencing costs per bp in relation to MiSeq | Limit of read length (usually up to a total of 300 bp), which would only fit small library sizes (up to ~260 bp) to ensuring overlapping paired-end reads |
| **Post-sequencing data treatment** | merging reads | reconstruction of entire haplotypes, including the original locus length; allows for global alignment among alleles. | If the total length of the reads is smaller than part of the library selected in the lab, larger fragments won't be merged and therefore not included in the analysis. |
| | *in-silico* size selection | Allow the selection of high coverage loci | Part of the sequences are filtered out because they are out of range |

### Supplementary Figures

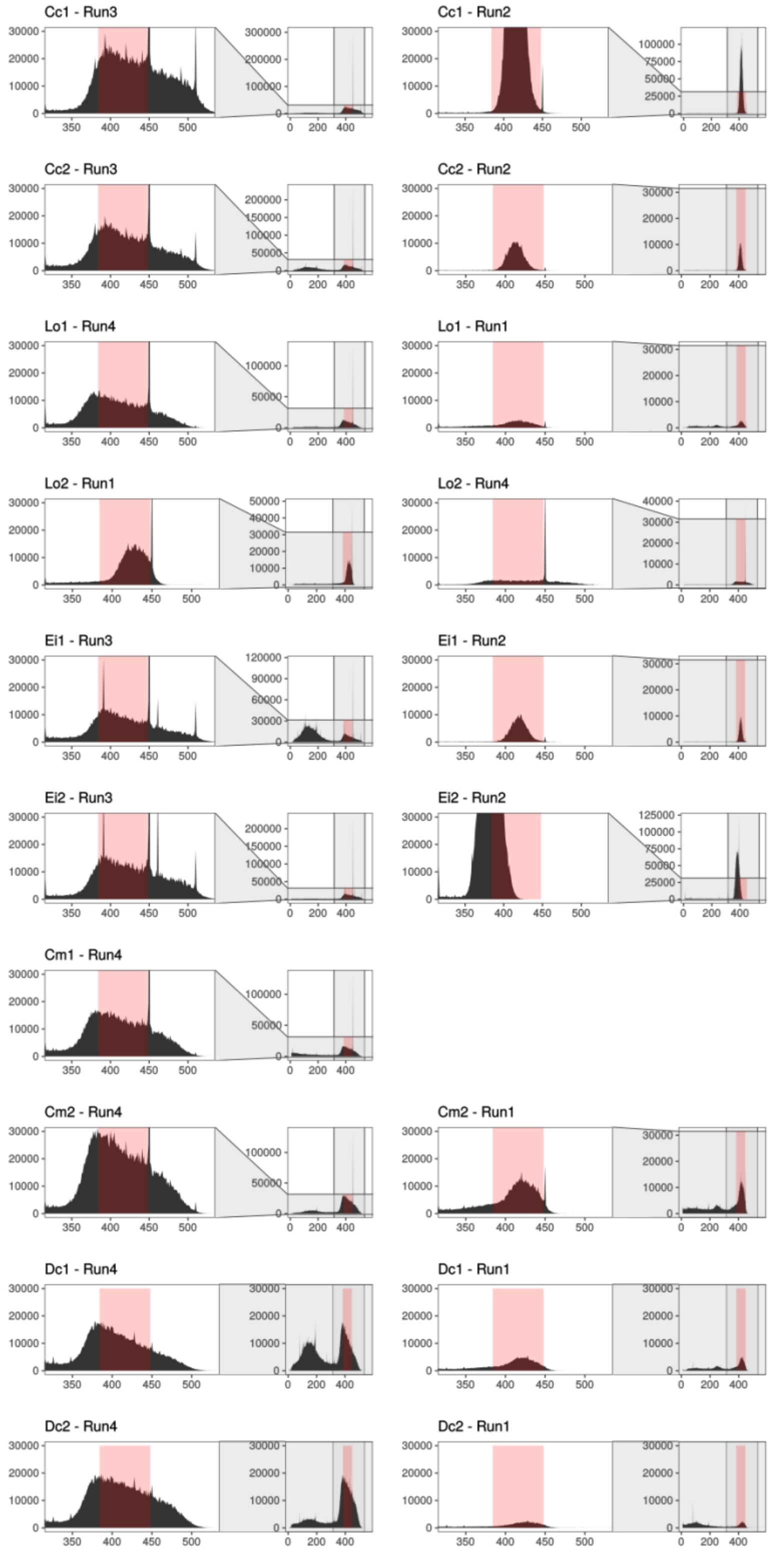


**Figure S1**. Fragment size distribution, of merged and preprocessed reads, for 10 sea turtle individuals (one per row) from 5 species across four sequencing runs. The zoomed-in area on the left side shows the main distribution area for each individual. In light red the in silico selected size range is highlighted. The left column shows all selected runs while the right column shows the non-selected runs.

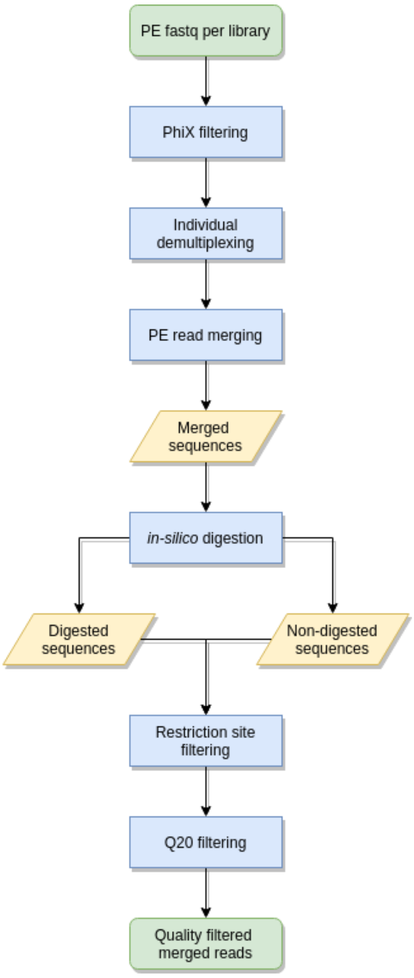


**Figure S2.** Read preprocessing workflow. PE: paired-end.

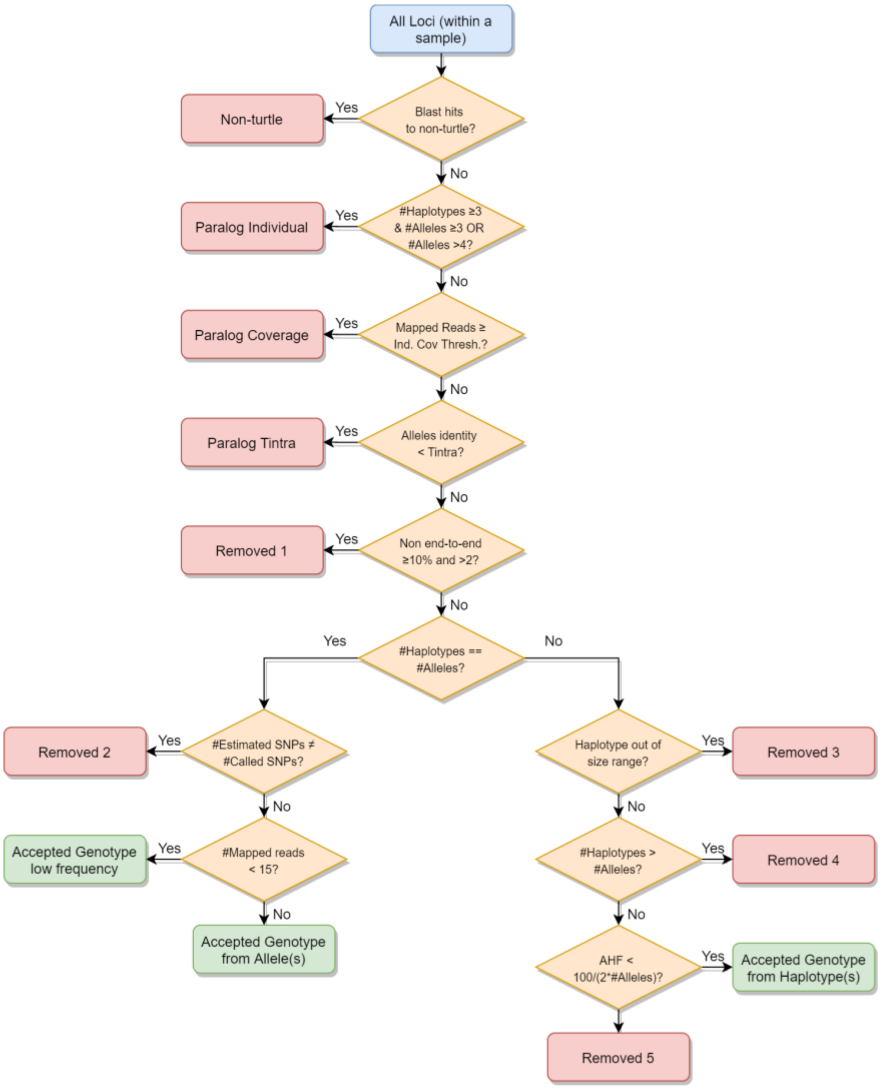


**Figure S3.** Decision tree for classification of loci clustered within individuals using the R2SCO pipeline. Acceptance of a locus depends on the reliability of the genotype called. The input data (blue square on top) include alleles from all loci defined after clustering above $T_{ances}$ and *sam* files including merged reads mapped to each locus. Orange rhombi represent each checking step, pink squares identify the non-turtle, paralogous and removed loci, and the green squares identify the accepted loci. The additional haplotype frequency (AHF) denotes the frequency of the first haplotype that exceeds the number of alleles when there are more haplotypes than alleles.

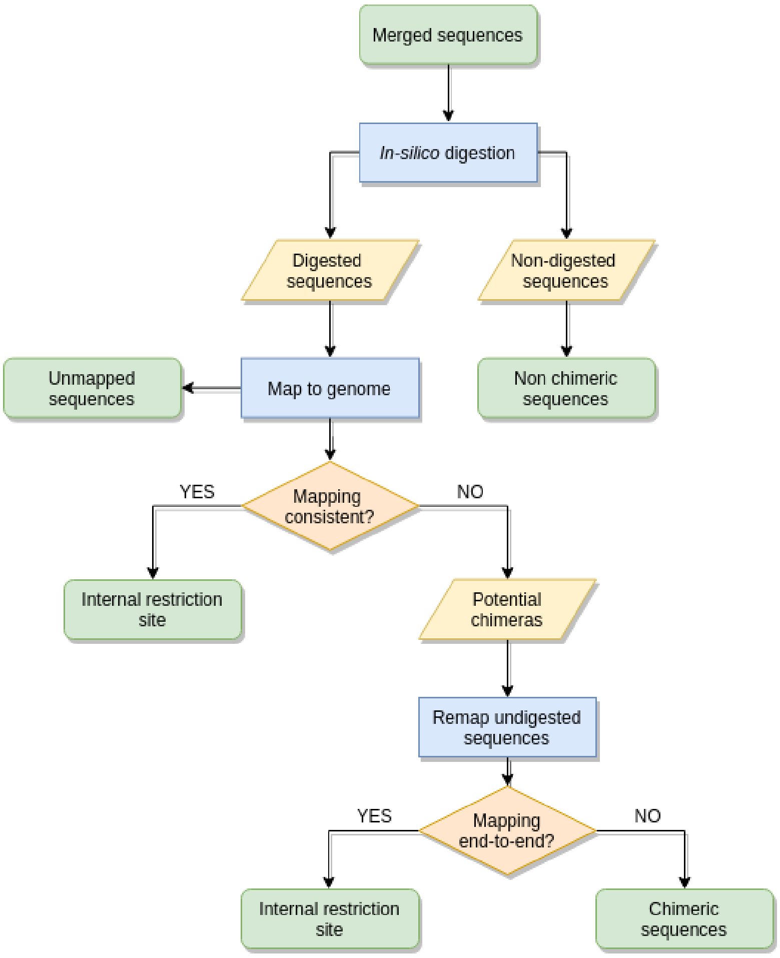


**Figure S4.** Workflow for chimera detection.

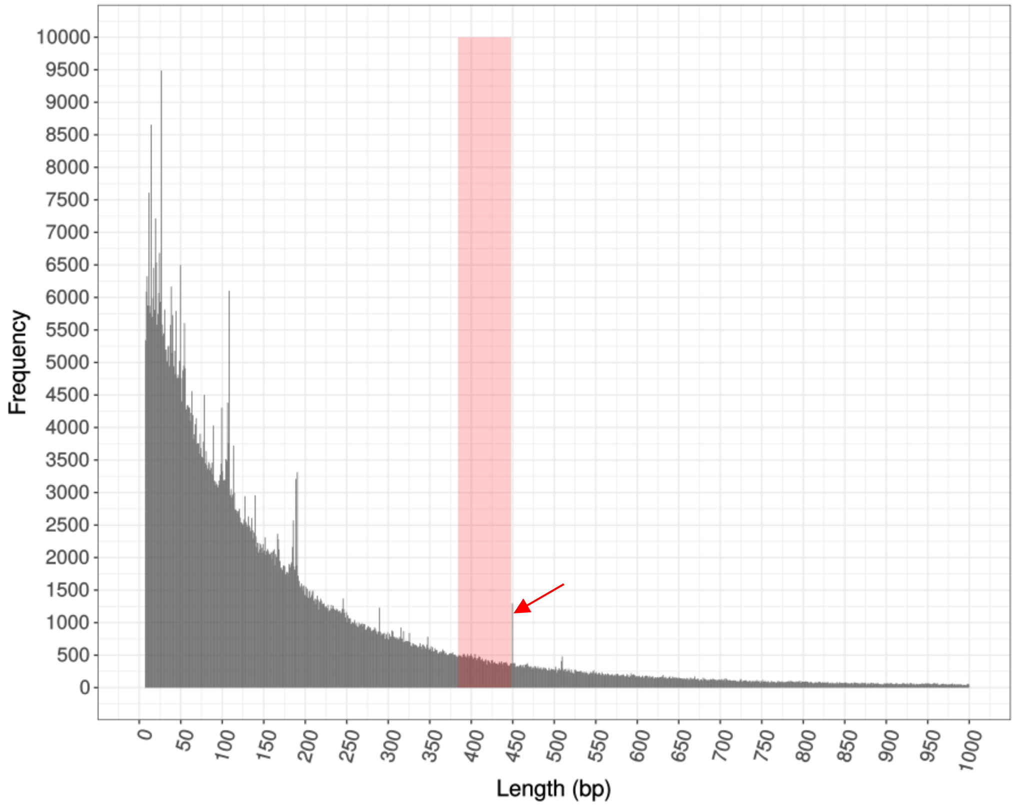


**Figure S5**. *C. mydas* genome *in silico* digestion including the *in silico* size range selected. The red arrow represents a disproportionately high peak representing paralogous loci in the genome.

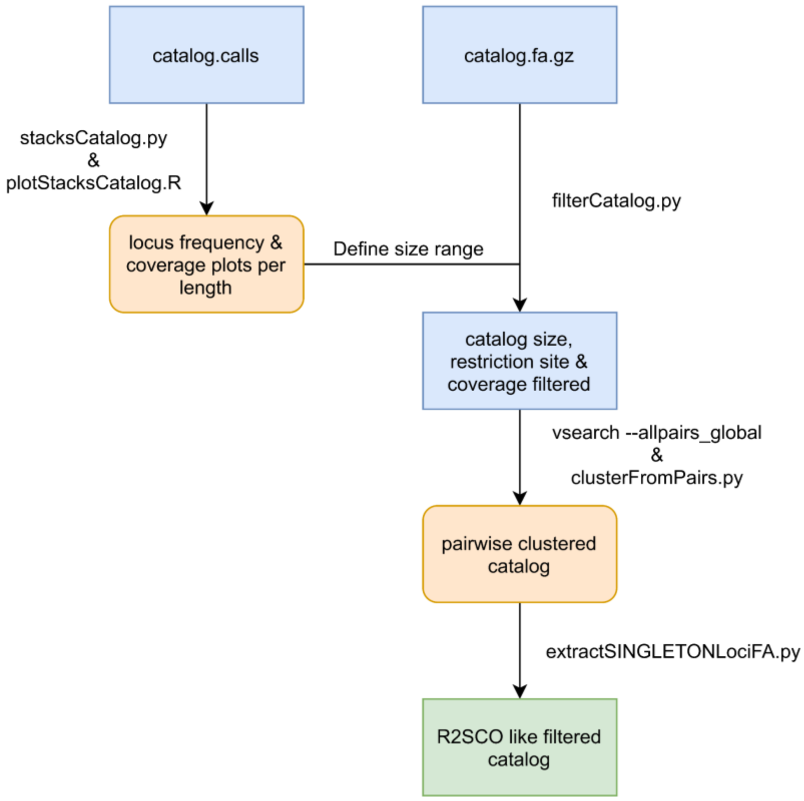

**Figure S6.** Stacks2R2SCOs workflow to filter the stacks catalog to generate an R2SCO like filtered catalog.

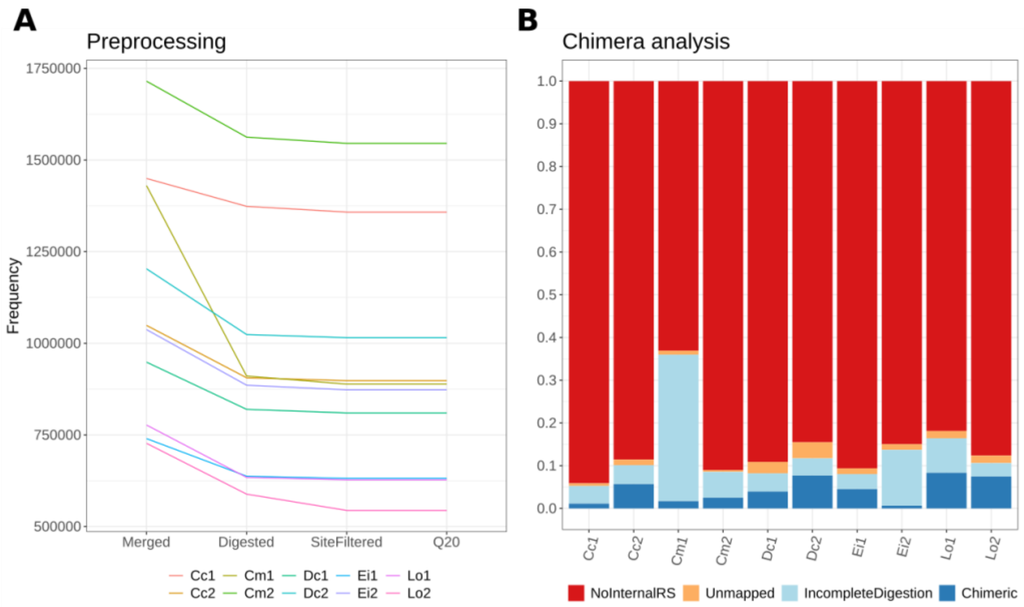


**Figure S7**. Results of preprocessing analysis within the selected size range (384-448 bp). (**A)** shows the preprocessing statistics for the ten different samples and (**B)** presents the percentage of digested fragments identified as either chimeras, with an incomplete digestion or inconclusive classification (i.e. unmapped). RS: restriction site; Q: Phred quality.

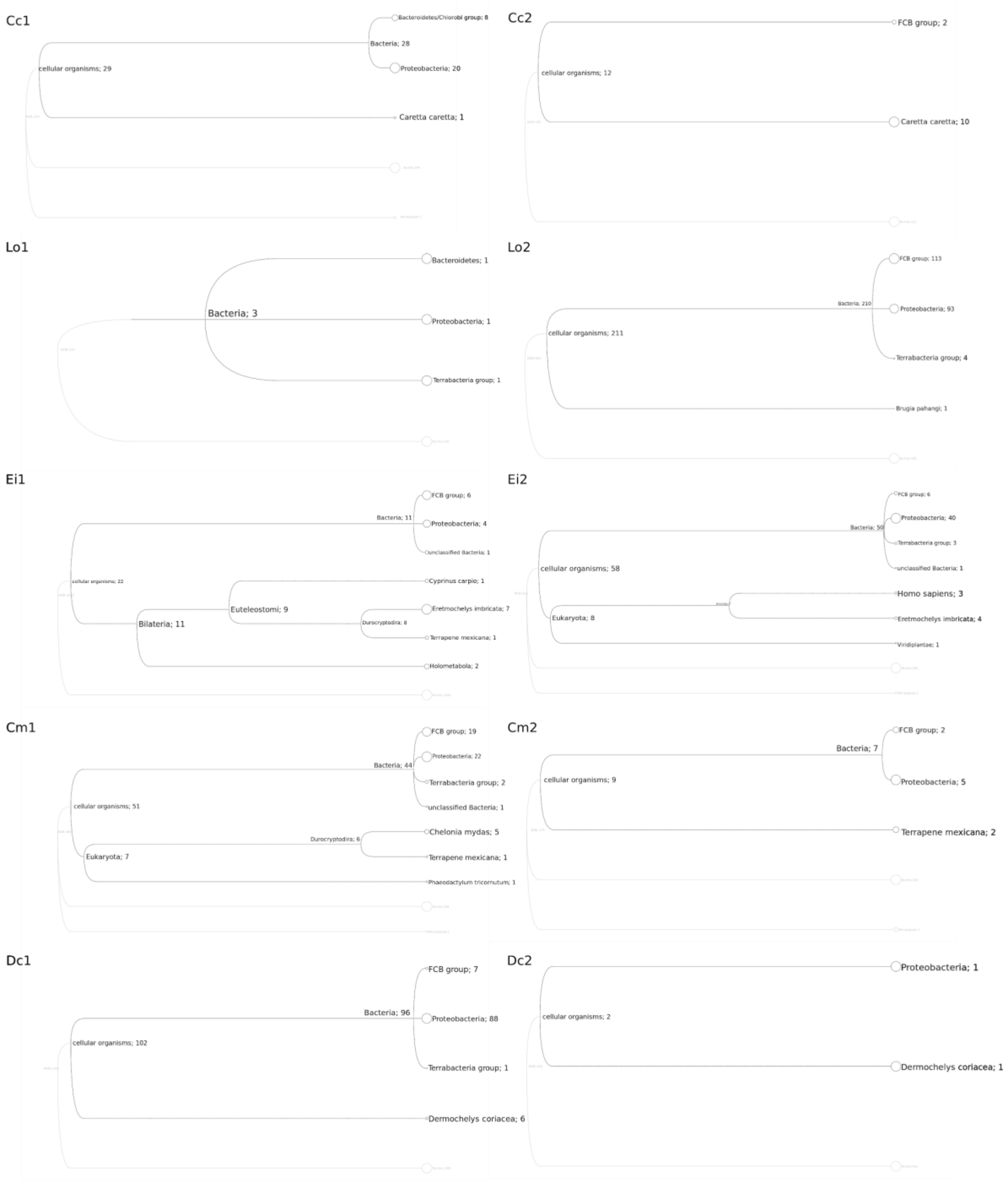


**Figure S8.** Megan trees from preliminary contamination analysis.

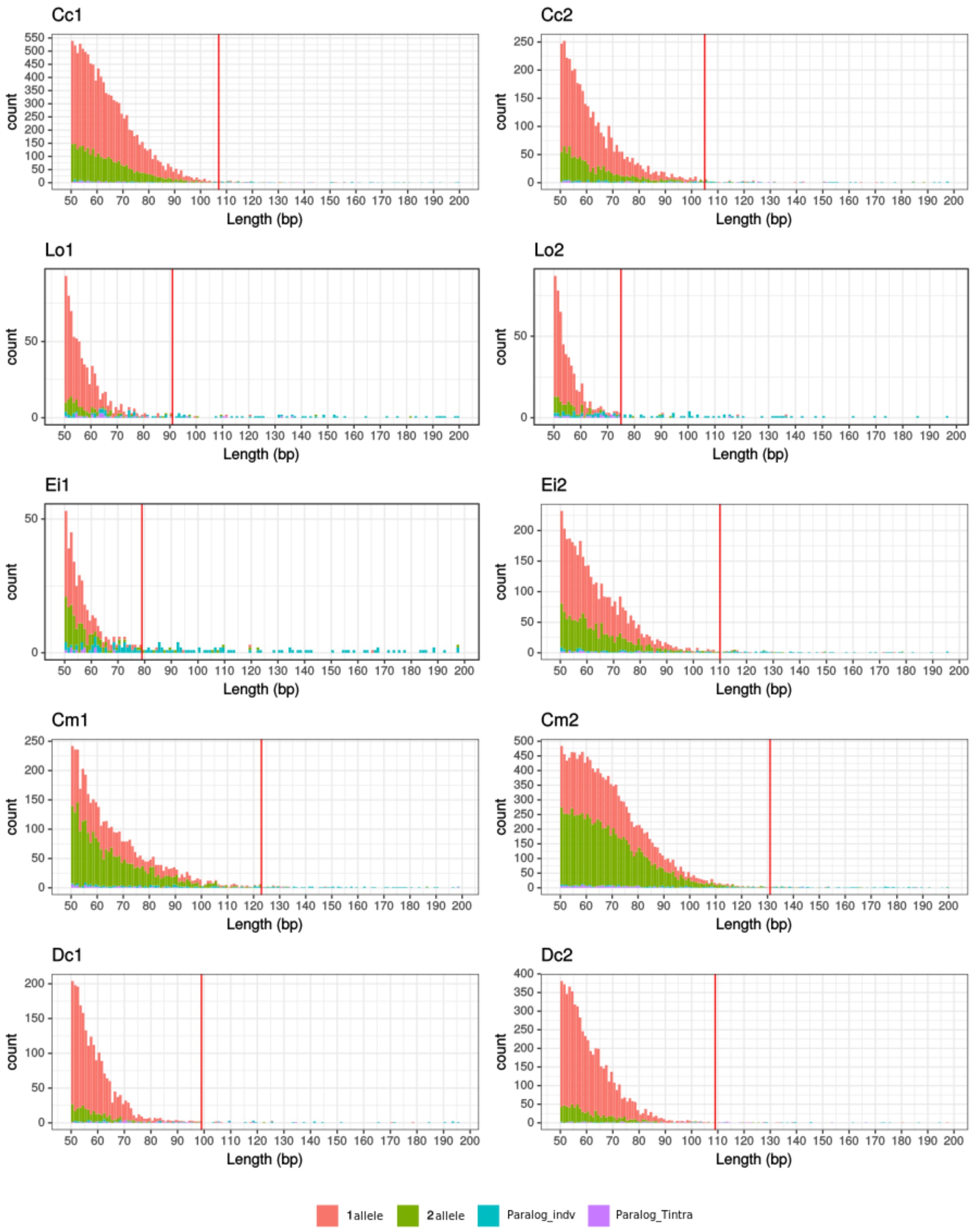


**Figure S9.** Coverage distribution across loci within each individual. The x axis starts at fragment size 50 bp. The red line indicates the $COV_{max}$ threshold set for each individual based on the end of distribution of coverages from putatively single-copy loci. 1 allele: clusters with a single allele; 2 alleles: clusters with 2 alleles; Paralog_indiv: clusters with more than 2 alleles; Paralog_Tintra: clusters containing two alleles with identity below the threshold $T_{intra}$.

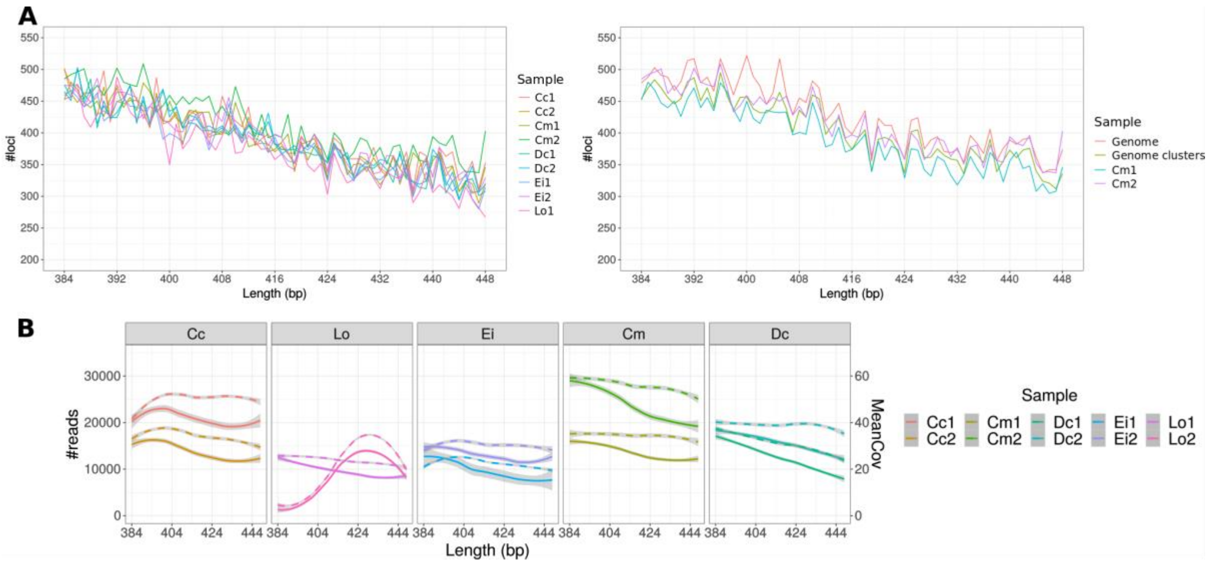


**Figure S10.** Number of loci, total coverage per fragment size and average coverage per locus across the selected size range for each individual. (**A**) Number of loci per fragment size estimated after clustering ddRAD data for 9 individuals and comparing the two *C. mydas* individuals (Cm1 and Cm2) and CheMyd_1.0_DNAzoo digested fragments before (Genome) and after (Genome clusters) clustering them using $T_{ances}$. The individual Lo2 was removed from this analysis as it presented very low coverage for the first part of the distribution. (**B**) Total amount of reads (y axis left, full line) vs. average locus coverage (y axis right, dotted line) per length (x axis) for each of the 10 individual selected runs. Lines were smoothed using local regression.

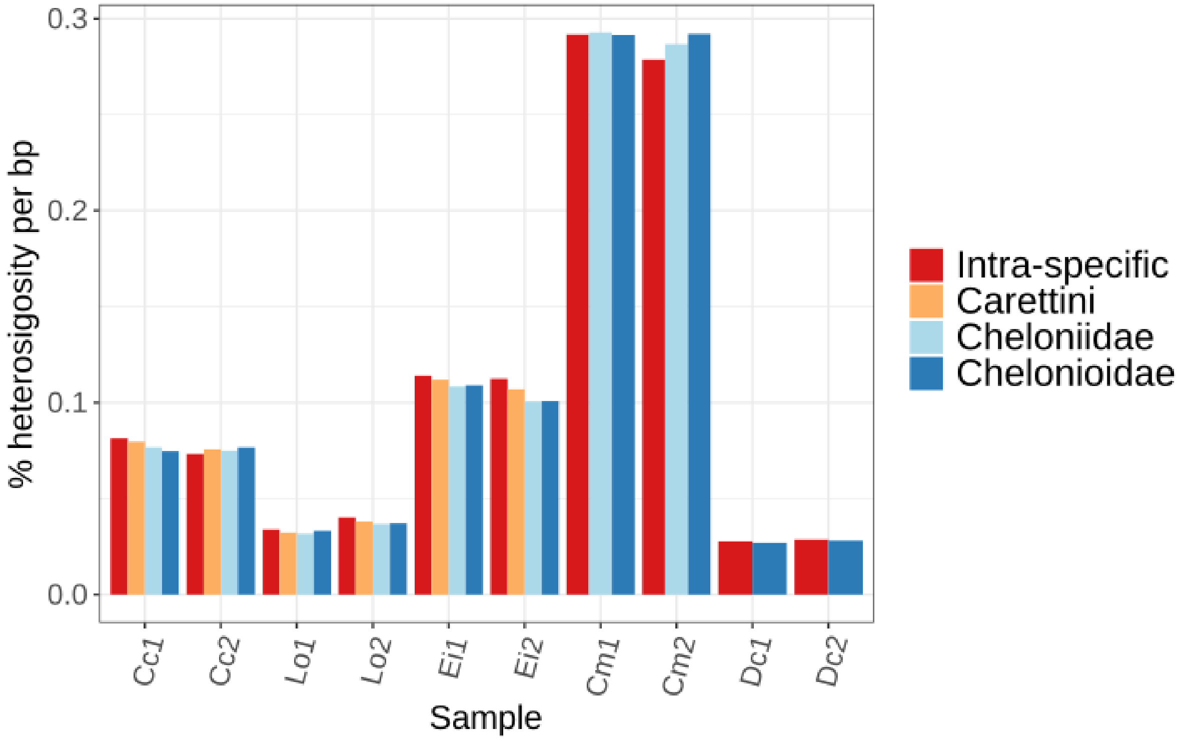


**Figure S11.** Intra-individual heterozygosity rates across different phylogenetic levels of R2SCOs.

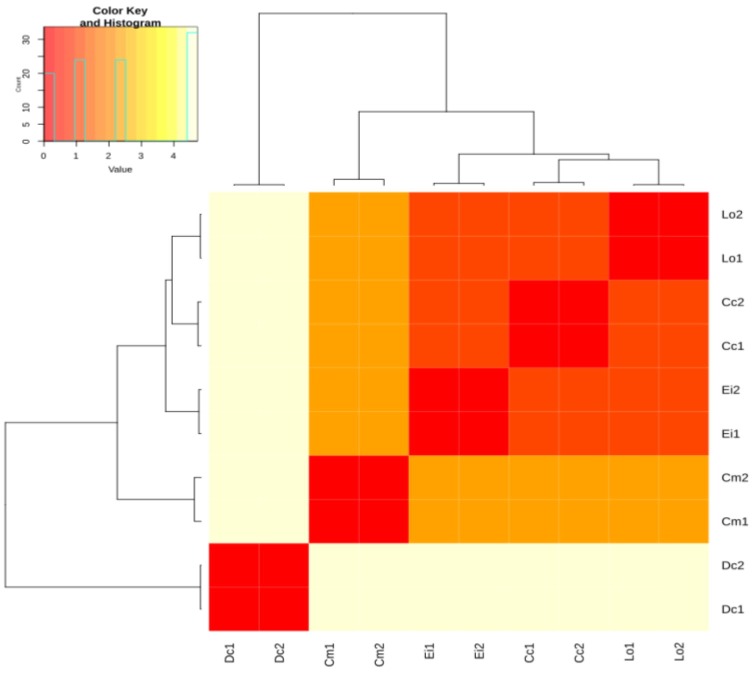


**Figure S12**. Heatmap of pairwise genetic distances across all individuals from the five sea turtle species. Genetic distance was calculated based on the total number of differences between the representative sequences of shared loci averaged by the total sequence length. The distance is highlighted by the gradient ranging from low distances (red) to larger distances (white/beige).

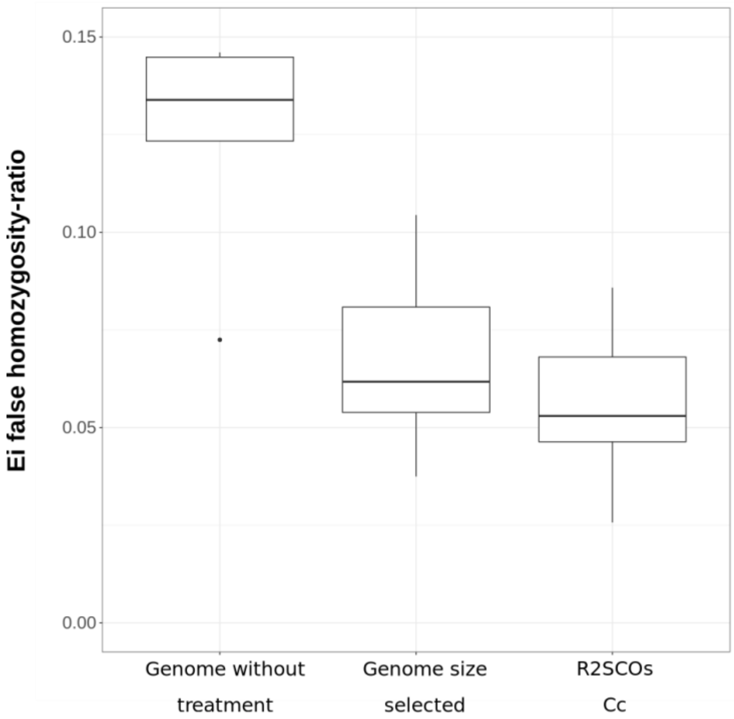


**Figure S13.** Ratio of incorrectly assigned genotypes using different references (*C. mydas* genome without treatment, *C. mydas* genome size selected and R2SCOs for *C. caretta* size selected) for backcrossed hybrids (*C. caretta* x *E. imbricata* hybrid) X *C. caretta* analyzed in Arantes et al., 2020b. R2SCO reference showed the lowest rate of false-homozygotes for *E. imbricata* (homozygotes for *E. imbricata* alleles are not expected in the backcrossed individuals) compared to the other references, although not at a significant level.

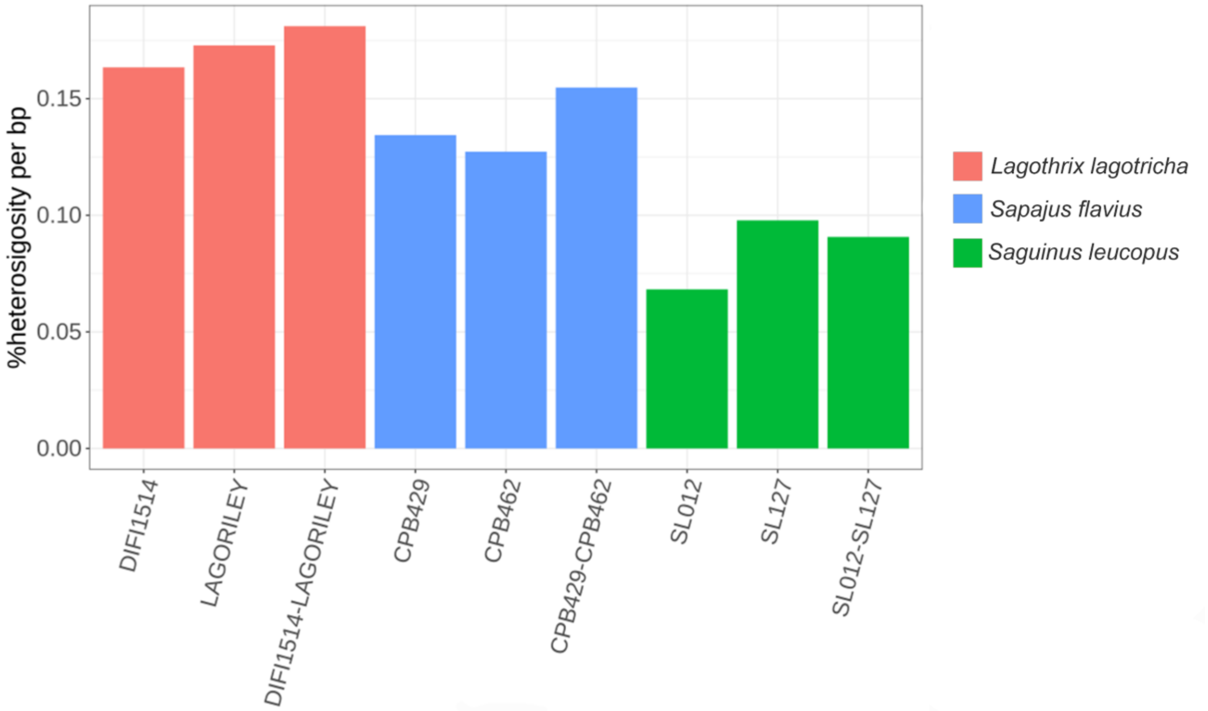


**Figure S14**. Heterozygosity levels for R2SCOs from primates from Valencia et al. 2018. The heterozygosity was estimated for each individual and genetic distances between individuals. The third bar for each species corresponds to the analysis of conspecific individuals.

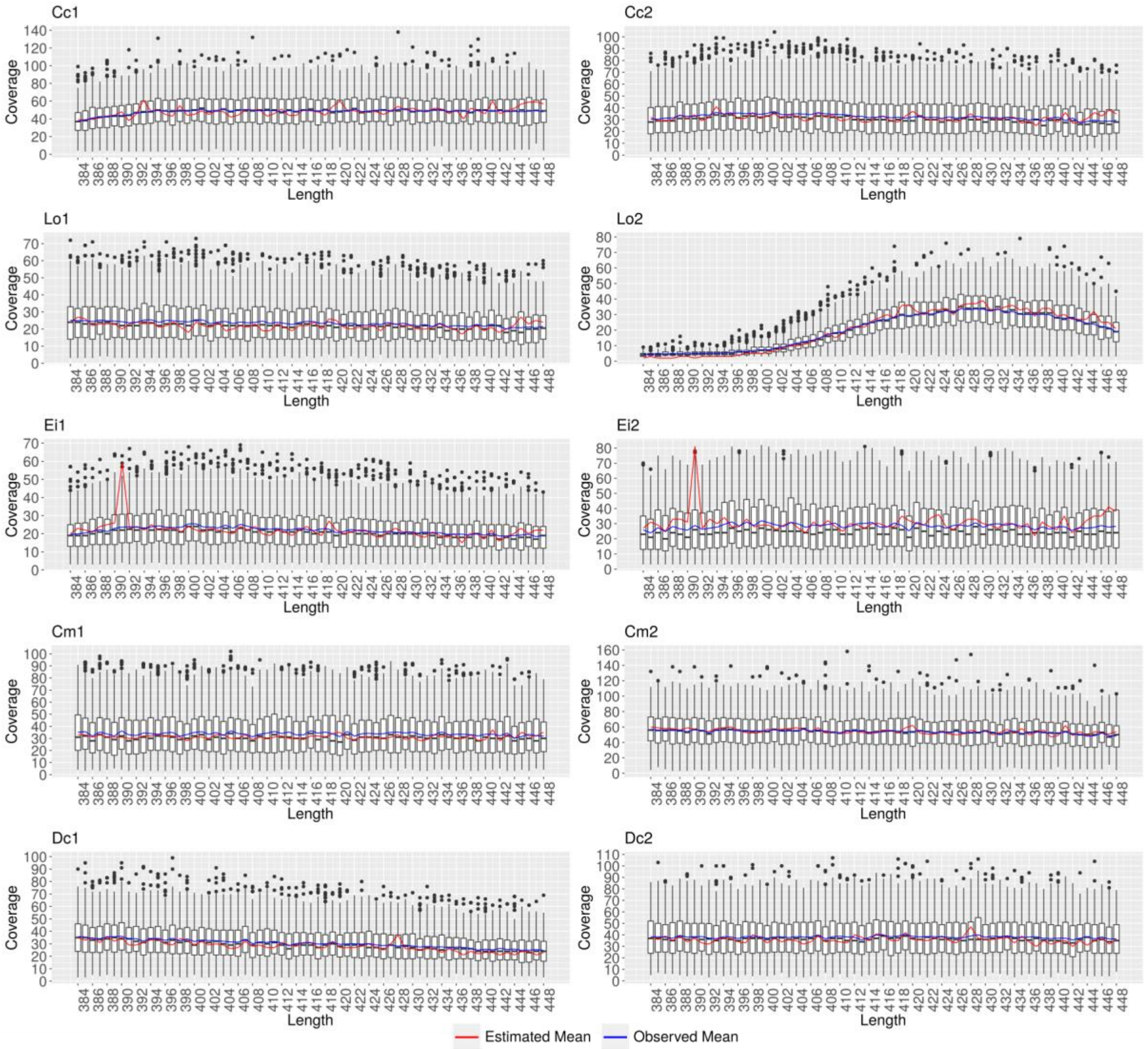


**Figure S15**. Boxplots representing the distribution of coverage per locus across fragment size. The blue line represents the observed average coverage found based on mapping reads to loci and the red line represents the estimated coverage based on the number of reads per fragment size divided by the number of fragments digested from the genome per size. The peak found around size 390 bp in Ei1 and Ei2 represents a peak in read coverage in the ddRAD data that is not shared by other species, including the genome.

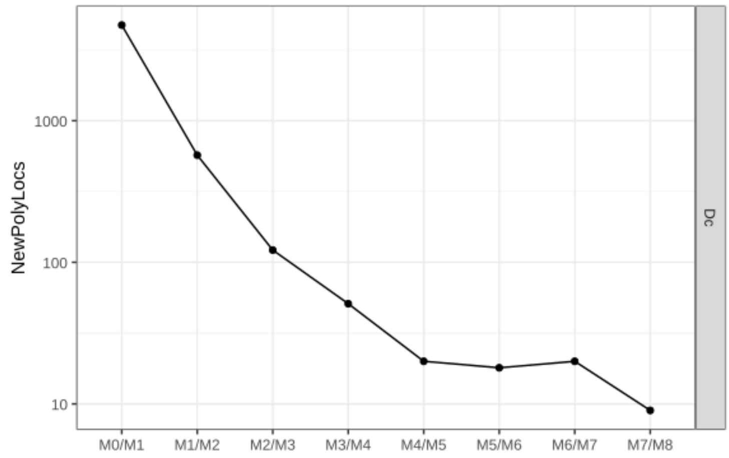


**Figure S16.** Difference in number of new polymorphic loci added for each iteration of *M* for *Dermochelys coriacea.*

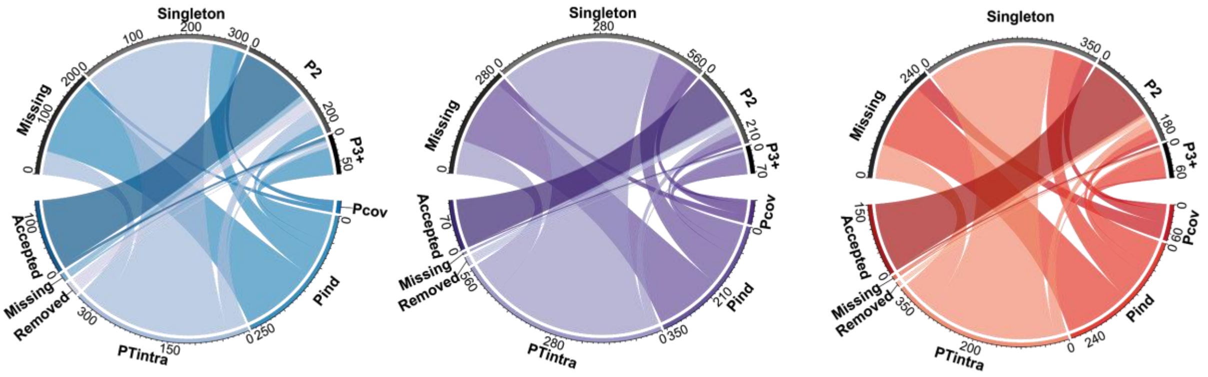


**Figure S17**. Circos plots showing comparisons of two *C. mydas* individuals (Cm1 on the left and Cm2 on the right) and their combined loci (Cm species in the middle) against the genome clusters of digested sequences. Only the putative loci relations are shown in detail.
